# Predicting the partition of behavioral variability in speed perception with naturalistic stimuli

**DOI:** 10.1101/601161

**Authors:** Benjamin M. Chin, Johannes Burge

## Abstract

A core goal of visual neuroscience is to predict human perceptual performance from natural signals. Performance in any natural task can be impacted by at least three sources of uncertainty: stimulus variability, internal noise, and sub-optimal computations. Determining the relative importance of these factors has been a focus of interest for decades, but most successes have been achieved with simple tasks and simple stimuli. Drawing quantitative links directly from natural signals to perceptual performance has proven a substantial challenge. Here, we develop an image-computable (pixels in, estimates out) Bayesian ideal observer that makes optimal use of the statistics relating image movies to speed. The optimal computations bear striking resemblance to descriptive models proposed to account for neural activity in area MT. We develop a model based on the ideal, stimulate it with naturalistic signals, predict the behavioral signatures of each performance-limiting factor, and test the predictions in an interlocking series of speed discrimination experiments. The critical experiment collects human responses to repeated presentations of each unique image movie. The model, highly constrained by the earlier experiments, tightly predicts human response consistency without free parameters. This result implies that human observers use near-optimal computations to estimate speed, and that human performance is near-exclusively limited by natural stimulus variability and internal noise. The results demonstrate that human performance can be predicted from a task-specific statistical analysis of naturalistic stimuli, show that image-computable ideal observer analysis can be generalized from simple to natural stimuli, and encourage similar analyses in other domains.

## Introduction

Human beings are remarkably adept at a wide variety of fundamental sensory-perceptual tasks. A sufficiently difficult task, however, can reveal the limits of human performance. A principal aim of perception science and systems neuroscience is to determine the limits of performance, and then to determine the sources of those limits. Performance limits and the factors that determine them have been rigorously investigated with simple tasks and stimuli. Ultimately, perception science aims to achieve the same rigorous understanding of how vision works in the real world. The current manuscript works towards this understanding by building upon recent advances for predicting the properties and performance of visual systems from natural stimuli.

In natural viewing, there exist at least three factors that limit performance: natural stimulus variability, internal noise, and suboptimal computations. Testing the relative importance of these sources requires two ingredients: i) an image-computable (images in, estimates out) ideal observer that specifies optimal performance in the task, and ii) experiments that can distinguish the behavioral signatures of each factor. Here, we develop theoretical and empirical methods that can predict and diagnose the impact of each source in mid-level visual tasks with natural and naturalistic stimuli. We apply these methods to the specific task of retinal speed estimation, a critical ability for estimating the motion of objects and the self through the environment.

When a pattern of light falls on the retina, millions of photoreceptors transmit information to the brain about the visual scene. The visual system uses this information to build stable representations of stimulus properties (i.e., latent variables) that are relevant for survival and reproduction. The visual system successfully extracts these critical latent variables from natural images in spite of tremendous stimulus variability; infinitely many unique retinal images (i.e. light patterns) are consistent with each value of a given latent variable. Some image features that vary across different natural images are particularly informative for extracting the latent variable(s) of interest. These are the features that the visual system should encode. Many other image features carry no relevant information. These features should be ignored. Variation in both the relevant and irrelevant feature spaces can limit performance. But the impact of stimulus variability on performance is minimized only if all relevant features are encoded. Thus, stimulus variability can differentially impact performance depending on the quality of feature encoding.

Signal detection theory posits that sensory-perceptual performance is based on the value of a decision variable^1^. But signal detection theory does not specify how to obtain the decision variable from the stimulus. Image-computable observer models do^2–6^. Image-computable *ideal* observer models specify how to encode and process the most useful stimulus features^7–13^. The explicit description of optimal processing provided by an image-computable ideal observer specifies how natural stimulus variability should propagate into the decision variable. Optimal processing minimizes stimulus-driven variation in the decision variable. Thus, stimulus variability and the optimal processing jointly set a fundamental limit on performance.

An image-computable ideal observer for estimating retinal image speed from local regions of natural images is shown in Fig. 1A. Given a set of stimuli, it uses the optimal computations (encoding receptive fields, pooling, decoding) for estimating speed from natural image movies^13^. The ideal observer therefore provides a principled benchmark against which to compare human performance. The tradition in ideal observer analysis is to constrain the ideal observer by stimulus and physiological factors known to limit the information content in the stimulus^9^. Natural stimulus variability and early measurement noise are two such factors that constrain the ideal observer (red text, Fig. 1A). The optimal computations govern how natural stimulus variability and early noise propagate into the ideal decision variable, thereby determining its variance (Fig. 1A). This variance is a critical determinant of ideal observer performance.

**Figure 1.**
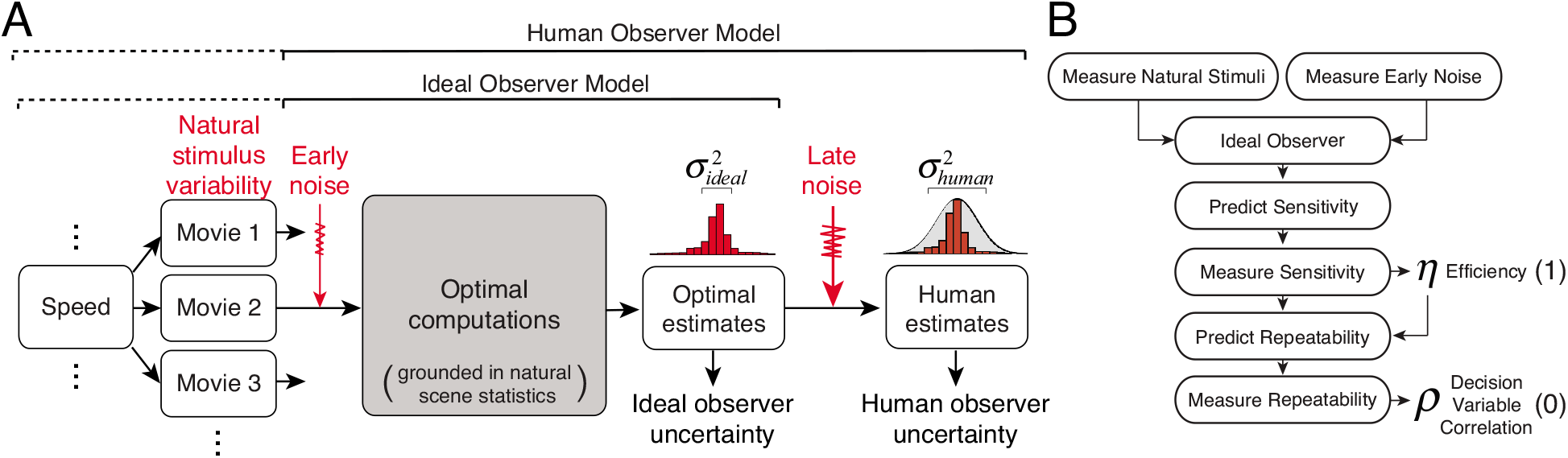
Ideal observer and plan for manuscript. **A** Ideal observer. Speed (i.e. the latent variable) can take on one of many values. Many different image movies share the same speed. The ideal observer is defined by the optimal computations (encoding, pooling, decoding) for estimating speed with natural stimuli. The optimal computations are grounded in natural scene statistics (gray box). For each unique movie, the ideal observer outputs a point estimate of speed. The ideal observer’s estimates vary across movies primarily because of natural stimulus variability, variability that is external to the observer. The degraded ideal observer is matched to overall human performance by adding late noise. **B** Plan for the manuscript. First, we measure natural stimuli and early noise to constrain an ideal observer for speed estimation. Next, we run an experiment and fit the efficiency of each human observer (1 free parameter) by comparing human to ideal sensitivity. Finally, we run a double-pass experiment and show that efficiency predicts human response repeatability (0 free parameters).

Human performance often tracks the pattern of ideal observer performance, but rarely achieves the same absolute performance levels. This is because humans are often limited by additional factors that are insufficiently well understood to include in the ideal observer. Ideal observers can therefore help identify and quantify the unknown factors responsible for the performance gap, and can serve as useful starting points for models of human performance.

To convert the ideal observer into a model of human observers, we consider two additional performance-limiting factors: late internal noise and suboptimal computations. Late internal noise is random, and is modeled at the level of the decision variable (Fig. 1A). Suboptimal computations (not depicted in Fig. 1A) are deterministic, and can be modeled at various computational stages. Both of these factors have the potential to increase the variance of the human decision variable compared to the ideal. Can the impact of these factors be distinguished experimentally, and what is the relative importance of each? Using complementary computational and experimental techniques, we show that in speed discrimination with naturalistic stimuli, i) humans underperform the ideal near-exclusively because of late internal noise, ii) the deterministic computations (encoding, pooling, decoding) performed by the human visual system are very nearly optimal, and iii) natural stimulus variability equivalently limits human and ideal performance. The work demonstrates that a task-specific analysis of naturalistic stimuli can tightly predict human performance, and shows that ideal observer analysis can be fruitfully applied to mid-level visual tasks with natural and naturalistic stimuli.

## Results

The plan for the manuscript is diagrammed in Fig. 1B. First, we develop an image-computable ideal observer model of retinal speed estimation that is constrained by measurements of natural stimulus variability and early noise, and then compare human to ideal performance in an experiment with matched stimuli. The first main experiment shows that humans track the predictions of the ideal but are consistently less sensitive: one free parameter—efficiency— accounts for the gap between human and ideal performance. Next, we hypothesize that human inefficiency is due to late internal noise, and not sub-optimal computations. This hypothesis predicts that natural stimulus variability should equally limit human and ideal observers. The second main experiment tests this hypothesis. Human observers viewed thousands of trials with natural stimuli in which each unique trial was presented twice. In this paradigm, the repeatability of responses reveals the respective roles of stimulus- and noise-driven variability. With zero additional free parameters, efficiency predicts response repeatability and the proportion of behavioral variability that is due to stimulus- and noise-driven components.

### Measuring natural stimuli

A fundamental problem of perception is that multiple proximal stimuli can arise from the same distal cause. This stimulus variability is an important source of uncertainty that will limit human and ideal speed discrimination performance. To measure natural stimulus variability, we photographed a large number of natural scenes^10,13^, and then drifted those photographs at known speeds behind a one degree aperture, approximately the size of foveal receptive fields in early visual cortex^14,15^. This procedure generates motion signals that are equivalent to those obtained by rotating the eye during smooth tracking of a target (Fig. 2A). The sampled set of stimuli approximates, but almost certainly underestimates, the variability present in the natural stimulus ensemble; looming and discontinuous motions, for example, are not represented in our training set^16,17^. Thus, the forthcoming estimates of the impact of naturalistic stimulus variability on ideal and human performance are likely to underestimate the impact of stimulus variability on human performance in natural viewing.

**Figure 2.**
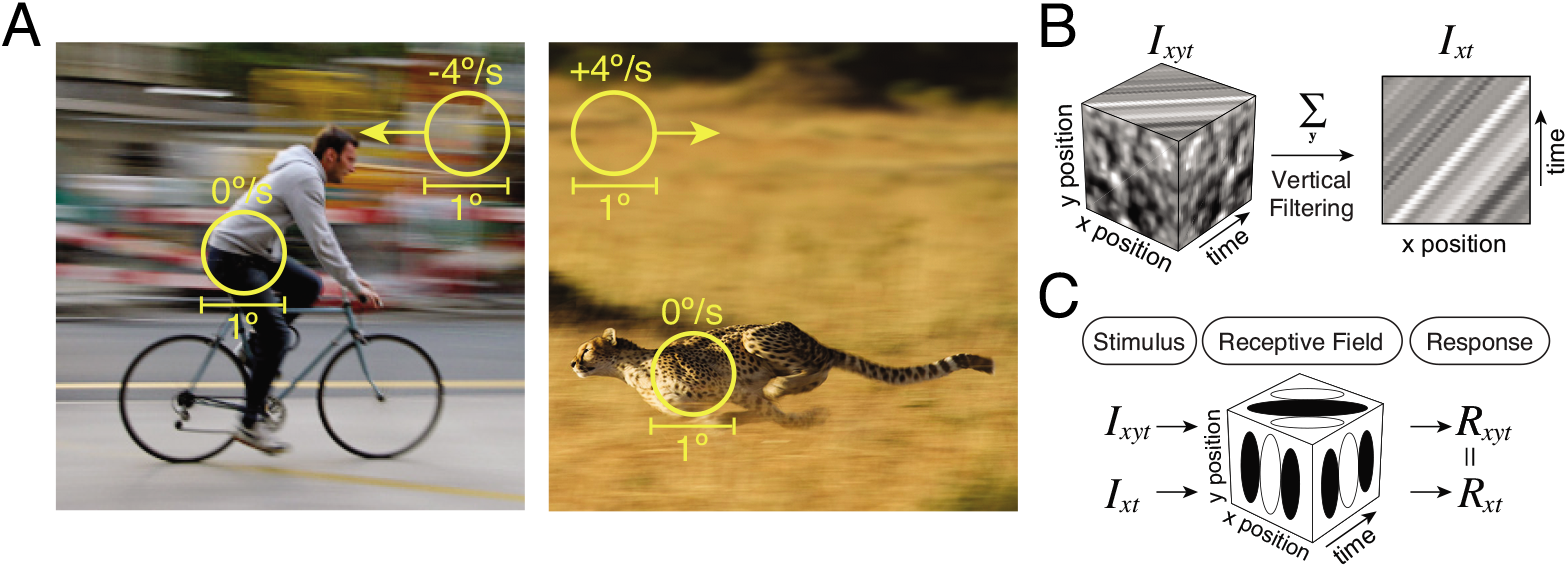
Naturalistic image movies and pre-processing. **A** Natural image movies were obtained by drifting photographs of natural scenes at known speeds behind one degree apertures for 250ms. Rotating the eye in its socket (e.g. tracking an object) creates the same pattern of motion in the stationary background. Optical properties of the eye and the temporal integration of the photoreceptors were also modeled. **B** Full space-time image movies (*Ixyt*) and vertically filtered space-time image movies (*Ixt*). Moving images can be represented as oriented signals in spacetime. **C** Vertically oriented receptive fields respond identically to full space-time movies and vertically filtered movies.

Movies drifted leftward or rightward with speeds ranging between 0.25 to 8.0 deg/sec. Movies were presented for 250ms, the approximate duration of a typical human fixation. The sampling procedure yielded tens of thousands of unique stimuli (i.e. image movies) at dozens of unique speeds. Image movies were then filtered so that only vertical orientations were present; that is, the stimuli were vertically averaged (i.e. *xt*) versions of full space-time (i.e. *xyt*) movies (Fig. 2B). Vertical averaging reduces stimulus complexity, but the resulting stimuli are still substantially more realistic than classic motion stimuli like drifiting sinewaves. Furthermore, vertically oriented receptive fields respond identically to vertically averaged and original movies (Fig. 2C). Thus, in an individual orientation column, the filtered movies should generate the same response statistics as the full space-time movies^13,18^. Finally, the contrast of the vertically-averaged stimuli were fixed to the modal contrast in natural scenes (see Discussion). Thus, our stimuli represent a compromise between simple and real-world stimuli, allowing us to run experiments with more natural stimuli without sacrificing quantitative rigor and interpretability. Our analysis should be generalizable to full space-time movies with more realistic forms of motion.

### Measuring early noise

All measurement devices are corrupted by measurement noise. The human visual system is no exception. Early measurement noise occurs at the level of the retinal image and places a fundamental limit on how well targets can be detected. Possible sources of early noise include the Poisson variability of light itself and the stochastic nature of the photoreceptor and ganglion cell responses^19^. The ideal observer for speed discrimination should be constrained by the same early noise as the human observer if it is to provide an accurate indication of the theoretically achievable human performance limits (see Fig. 1A).

Human observers performed a target detection task using the equivalent input noise paradigm^7,20^. The task was to detect a known target embedded in dynamic Gaussian white noise. On each trial, human observers viewed two stimuli in rapid succession, and tried to identify the stimulus containing the target (Fig. 3A,B). The time-course of stimulus presentation was identical to the forthcoming speed discrimination experiment. Figure 2C shows psychometric functions for target detection in one human observer as a function of target contrast. Each function corresponds to a different noise contrast. Detection thresholds, which are the target contrasts required to identify the target interval 76% of the time (i.e. d-prime of 1.0 in a 2IFC task), are shown for two different targets (3.0 and 4.5 cpd) in Fig. 3D. Consistent with previous studies, contrast power at threshold increases linearly with pixel noise^7,21^. Figure 2E shows the same data plotted on logarithmic axes, a common convention in the literature. There are two critical points on this function. The first is its value when pixel noise equals zero, where detection performance is limited only by internal noise. The second is at double the contrast power of the first point—the so-called ‘knee’ of the function—where the pixel noise equals the internal noise (see Supplement). This level of pixel noise is known as the equivalent input noise.

**Figure 3.**
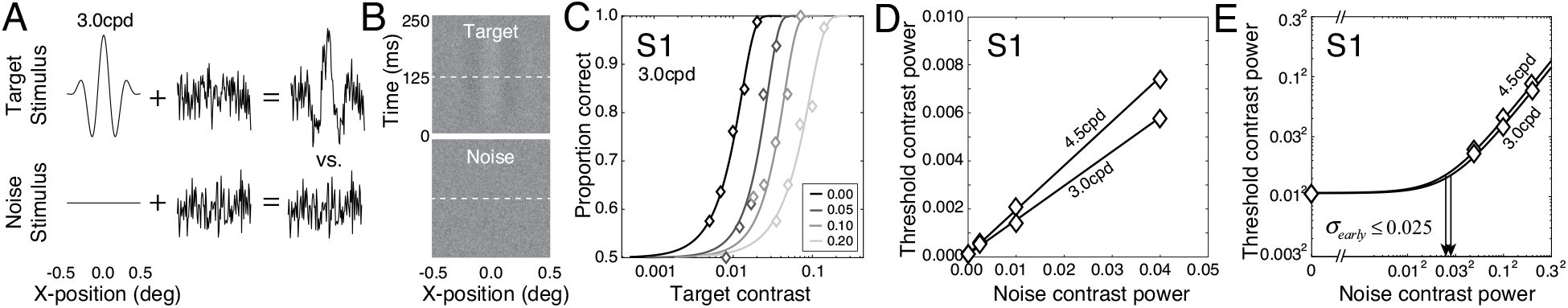
Measuring early noise with a target detection experiment. **A** Stimulus construction. On each interval, the stimulus was either a target sinewave or a middle gray luminance pattern corrupted by dynamic noise. **B** On each trial, the task was to report which of two intervals contained the target stimulus. **C** Psychometric functions from one human observer (S1) for different noise contrasts. **D** Target contrast power at the detection threshold for the same human observer. Thresholds increase linearly with noise contrast power. **E** Target contrast power at detection threshold plotted on a log-log axis (same data as D). Arrows indicate the estimate of equivalent input noise.

Internal noise was estimated separately for each target type and human observer. Estimates were consistent across target types and were thus averaged. Internal noise estimates for the first, second, and third human observers are 2.5%, 2.3% and 2.9%, respectively (Fig. S1). These values are in line with previous reports^7,20,22^. It has been argued that the controlling internal noise in target detection experiments may be due only to early noise^21^. This argument is controversial. The controlling noise could be early, but it could also arise at later (e.g. decision) stages. The equivalent input noise thus represents an upper bound on the amount of early noise in the human visual system. Because the upper bound is small, plausible amounts of early noise within the bound only weakly impact ideal observer performance (see below, Fig. S2). The ideal observer that we present in the main text is limited by early noise at this upper bound.

### Ideal observer

An ideal observer performs a task optimally, making the best possible use of the available information given stimulus variability and specified biological constraints. In addition to natural stimulus variability and early noise (see Figs. 2,3), we model the optics of the eye^23^, the temporal integration of photoreceptors^24^, and the linear filtering^25^ and response normalization^26–28^ of cortical receptive fields, because they are well established features of early visual processing and because they determine the information available for processing.

Assuming the relevant factors have been accurately modeled, ideal observers provide principled benchmarks against which to compare human performance. Humans often track the pattern but fail to achieve the absolute limits of ideal performance. As a consequence, ideal observers often serve as principled starting points for determining additional unknown factors that cause humans to fall short of theoretically achievable performance limits.

Developing an ideal observer with natural stimuli is challenging because it is unclear a priori which stimulus features are most useful for the task. We find the optimal receptive fields for speed estimation using a recently developed Bayesian statistical learning method called Accuracy Maximization Analysis^18,29,30^ (AMA). Given a stimulus set, the method learns the receptive fields that encode the most useful stimulus features for the task (Fig. S3A). Once the optimal features are determined, the next step is to determine how to optimally pool and decode the responses **R** = [*R*_1_,*R*_2_,⋯,*R_n_*] of the receptive fields that select for those features where *n* is the total number of receptive fields. Eight receptive fields capture essentially all of the useful stimulus information; additional receptive fields provide negligible improvements in performance^13^.

The optimal pooling rules are specified by the joint statistics between the receptive field responses and the latent variable^18,31^. With appropriate response normalization, the responses across stimuli for each speed are conditionally Gaussian^13,32,33^ (Fig. S3B). To obtain the likelihood of a particular speed, the Gaussian statistics require that the receptive field responses to a given stimulus be pooled via weighted quadratic summation (see Supplement; Fig. S3C). The computations for computing the likelihood thus instantiate an enhanced version of the motion-energy model, indicating that energy-model-like computations are the normative computations supporting speed estimation with natural stimuli^2,18^. The speed tuning curves of hypothetical neurons implementing these computations mimic the properties of speed tuning curves in area MT^34^ (Fig. SD). Finally, an appropriate read out of the population response of these hypothetical neurons is equivalent to decoding the optimal estimate from the posterior probability distribution *p*(*X*|**R**) over speed (Fig. S3EF). If a 0,1 cost function is assumed, the latent variable value corresponding to the posterior max is the optimal estimate. We have previously verified that reasonable changes to the prior and cost function do not appreciably alter the optimal receptive fields, pooling rules, and estimation performance^30^.

The factors thus far described in the paper—stimulus variability and early noise, biological constraints, and the optimal computations (encoding, pooling, decoding)—all impact ideal performance in our task. The precise amount of early noise is the only factor subject to some uncertainty, given a stimulus set. However, within the bound set by the detection experiment (see Fig. 3), different amounts of early noise have only a minor effect on ideal performance (Fig. S2). Thus, estimates of ideal performance are stable and set overwhelmingly by stimulus variability.

### Measuring efficiency

The ideal observer benchmarks how well humans use the stimulus information available for the task. Efficiency quantifies how human sensitivity *d*′_*human*_ compares to ideal observer sensitivity *d*′_*ideal*_ and is given by

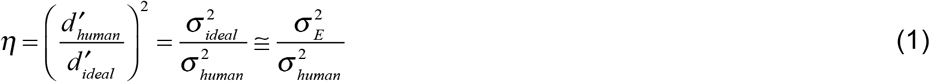

where 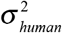 is the total variance of the human decision variable, 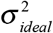 is the total variance of the ideal decision variable, and 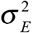 is the stimulus-driven component of the ideal decision variable. The third approximate equality in Eq. 1 assumes that stimulus-driven variability equals ideal observer variability because the impact of early noise is bounded to be small (see Fig. 3). Supplementary analyses show that the approximation and the uncertainty about the precise amount of early noise within the bound do not affect estimates of human efficiency by more than 10% (see Methods; Fig. S2).

To measure human sensitivity, we ran a two-interval forced choice (2IFC) speed discrimination experiment. On each trial, human observers viewed two moving stimuli in rapid succession, and indicated which stimulus was moving more quickly (Fig. 4A). This design is similar to classic psychophysical experiments with one critical difference. Rather than presenting the same (or very similar) stimuli in each condition hundreds of times, we present hundreds of unique stimuli one time each. This stimulus variability jointly limits human and ideal performance. Human sensitivity is computed using standard expressions from signal detection theory 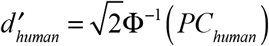 where *PC_human_* is the proportion of times that the comparison is chosen in a given condition in a 2IFC experiment and Φ^−1^(·) is the inverse cumulative normal. (This expression is correct assuming the observer uses the optimal criterion, an assumption that is justified by the data; Fig. S4)

**Figure 4.**
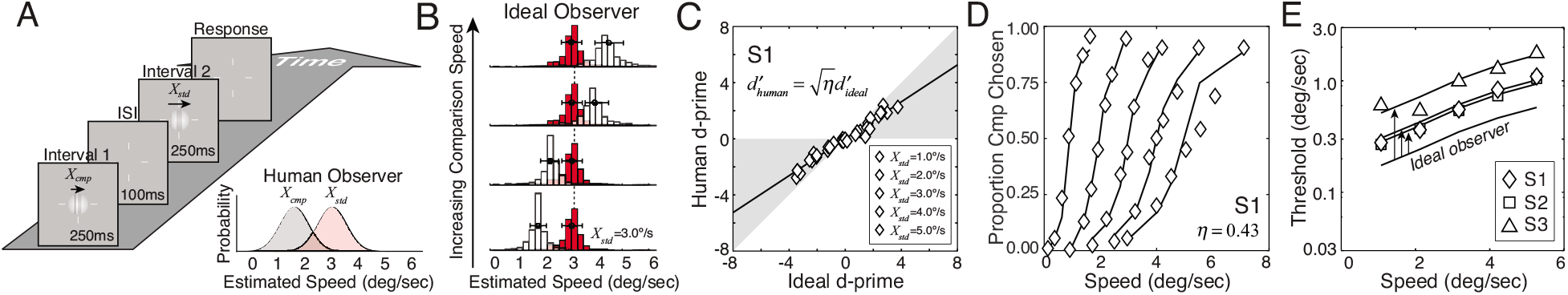
Measuring speed discrimination. **A** The task in a two-interval forced choice experiment was to report the interval containing the faster of two natural image movies. Unlike classic psychophysical studies, which present the same stimuli hundreds of times, the current study presents hundreds of unique stimuli one time each. This design injects naturalistic stimulus variability into the experiment. Human responses are assumed to be based on samples from decision variable distributions (inset). **B** Ideal observer estimates across hundreds of standard (red) and comparison movies (white) at one standard speed (3 deg/sec) and four comparison speeds. **C** Human vs. ideal observer sensitivity for all standard and comparison speeds. Shaded regions mark regions of plot where humans are less efficient than ideal but are still performing the task. For all conditions, humans are less sensitive than the ideal observer by a single scale factor: efficiency: 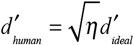 Negative d-primes correspond to conditions in which the comparison was slower than the standard. **D** Psychometric functions for one human observer (symbols) at five standard speeds. The degraded ideal observer (solid curves) matches the efficiency of the human observer (one parameter fit to human data). **E** Human speed discrimination thresholds (d-prime = 1.0) as a function of standard speed for three human observers (symbols) on a semi-log plot. The pattern of human thresholds matches ideal observer thresholds (solid curve). Vertically shifting the ideal observer thresholds by an amount set by each human’s efficiency (arrows) shows degraded observer performance (solid curves, one free parameter fit per human).

To measure ideal sensitivity, we ran the ideal observer in a simulated experiment with the same stimuli as the human. Ideal sensitivity (i.e. d-prime) was computed directly from the distributions of ideal observer speed estimates in each condition (Fig. 4B). Human and ideal sensitivities across all speeds are linearly related (Fig. 4C). Rearranging Eq. 1 shows that human sensitivity 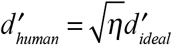 equals the ideal observer sensitivity degraded (scaled) by the square root of the efficiency. Thus, a single free parameter (efficiency) relates the pattern of human and ideal sensitivities for all conditions. The efficiencies of the first, second, and third human observers are 0.43, 0.41, and 0.17, respectively (Fig. S5).

Transforming the sensitivity data back into percent comparison chosen shows that the details of the degraded ideal nicely account for the human psychometric functions (Fig. 4D). The psychometric functions can be summarized by the speed discrimination thresholds (d-prime = 1.0; 76% correct in a 2IFC task). The pattern of human and ideal thresholds match; the proportional increases of the human and ideal threshold functions with speed are the same (Fig. 4E). These results quantify human uncertainty 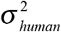, show that an ideal observer analysis of naturalistic stimuli predicts the pattern of human speed discrimination performance, and replicate our own previously published findings^13^.

Together, the ideal observer and speed discrimination experiment reveal the degree of human inefficiency (i.e. how far human performance falls short of the theoretical ideal). But they cannot determine the sources of this inefficiency. Humans could be inefficient because of late noise. Humans could also be inefficient because of suboptimal computations. If inefficiency is due exclusively to late noise, stimulus variability must equally limit human and ideal observer performance. If human inefficiency is partly due to suboptimal computations, stimulus variability will cause more stimulus-driven uncertainty in the human than in the ideal. How can human behavioral variability be partitioned to determine the sources of inefficiency in speed perception? To do so, additional experimental tools are required.

### Predicting and measuring decision variable correlation

A double pass experiment, when paired with ideal observer analysis, can determine why human performance falls short of the theoretical ideal. In a double pass experiment^35–37^, each human observer gives a response to each of a large number of unique trials (i.e. the first pass), and then performs the entire experiment again (i.e. the second pass). Double pass experiments can ‘unpack’ each point on the psychometric function (Fig. 5AB), providing far more information about the factors driving and limiting human performance than standard single pass experiments. The correlation in the human decision variable across passes—decision variable correlation—is key for identifying the factors that limit performance and determine efficiency^35,38^.

**Figure 5.**
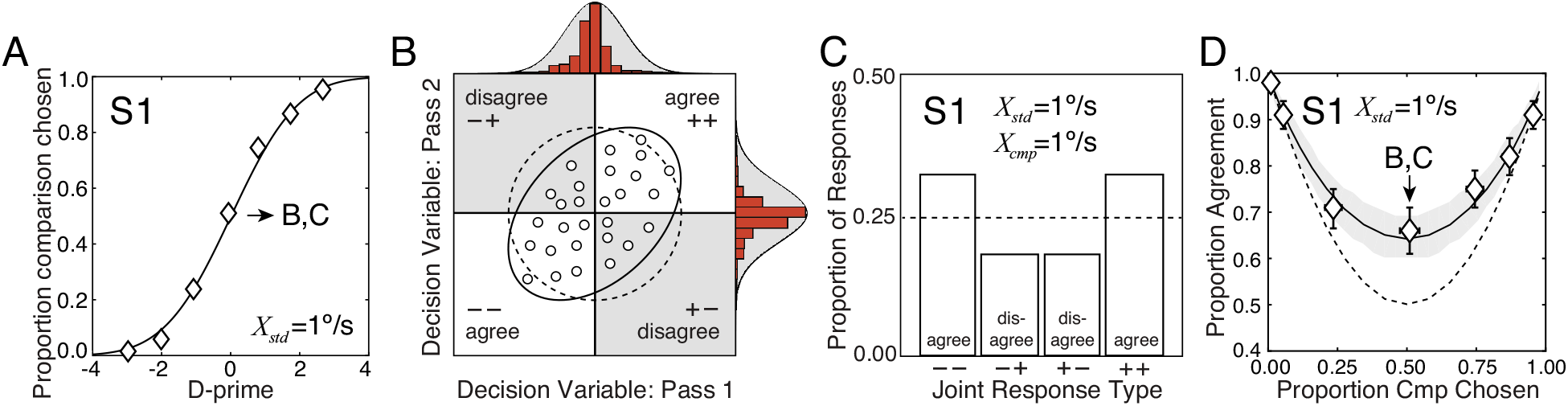
Decision variable correlation and response repeatability in a double pass experiment. **A** Psychometric data from the first human observer and cumulative Gaussian fit plotted as proportion comparison chosen vs. d-prime for the standard speed of 1 deg/sec. (Same data as in Fig. 4D.) **B** Schematic for visualizing decision variable correlation across passes when standard and comparison speeds are identical (e.g. both equal 1 deg/sec). Samples correspond to individual double pass trials (small circles). The value of each sample represents the difference between the estimated speeds of the comparison and standard stimuli on each trial. Decision variable values corresponding to response agreements and disagreements fall in white and gray quadrants, respectively. Decision variable distributions with the decision variable correlation predicted by efficiency (solid ellipse) and by the null model with a decision variable correlation of zero (dashed ellipse). Decision variable correlation depends on the relative importance of correlated and uncorrelated factors across passes. Natural stimulus variability is correlated on each repeated trial of a double pass experiment; internal noise is not. Criteria on each pass (vertical and horizontal lines, respectively) are assumed to be optimal and at zero. **C** Predicted response counts (bars) for each response type (−−, −+, +−, ++) across passes (100 trials per condition) given the decision variable correlation shown in B. **D** Proportion of trials on which responses agreed across both passes of the double pass experiment as a function of proportion comparison chosen for one human observer. Agreement data (symbols) and prediction (solid curve) assuming that efficiency predicts decision variable correlation (i.e. that all human inefficiency is due to late noise). The null prediction assumes that the decision variable correlation across passes is zero (dashed curve). The agreement data is predicted directly from the efficiency of the human observer (zero free parameters). Error bars represent 68% bootstrapped confidence intervals on human agreement. Shaded regions represent 68% confidence intervals from 10000 Monte Carlo simulations of the predicted agreement data assuming 100 trials per condition.

The power of this experimental design is that it enables behavioral variability to be partitioned into correlated and uncorrelated factors. Factors that are correlated across passes, like the stimuli, increase the correlation of the decision variable across passes. Factors that are uncorrelated across passes, like internal noise, decrease the correlation of the decision variable across passes. If the variance of the human decision variable is dictated only by stimulus-driven variability, decision variable correlation will equal 1.0. If the variance of the human decision variable is dictated only by internal noise, decision variable correlation will equal 0.0. If both stimulus-driven variability and internal noise play a role, the correlation will have an intermediate value.

Decision variable correlation, like the decision variable itself, cannot be measured directly using standard psychophysical methods. Rather, it must be inferred from the repeatability of responses across passes in each condition. The higher the decision variable correlation, the greater the proportion of times responses agree (i.e. repeat) in a given condition (Fig. 5BC; Fig. S6).

In each condition, we used the pattern of response agreement to estimate decision variable correlation (Fig. 5BC), and then plotted agreement against the proportion of times the human observer (symbols) chose the comparison stimulus as faster (Fig. 5D). Human response agreement implies a decision variable correlation that is significantly different from zero. For the seven conditions shown in Fig. 5D (i.e. all comparison speeds at the 1 deg/sec standard speed), the maximum likelihood fit of decision variable correlation across the seven comparison levels is 0.43. Thus, 43% of the total variance in the human decision variable is due to factors that are correlated across repeated presentations of the same trials.

How should the estimate of decision variable correlation be interpreted? Human decision variable correlation across passes is given by

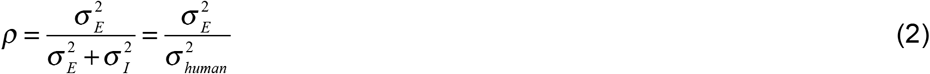

where 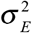 is the stimulus-driven variance, 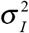 is the noise-driven variance, and 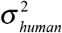 is the total variance of the human decision variable. Decision variable correlation is driven by stimulus variation, because the stimuli are perfectly correlated across passes.

The estimated decision variable correlation is strikingly similar to the efficiency measured for each observer. Although the exact relationship between decision variable correlation and efficiency depends on the source of human inefficiency, the fact that they are similar is no accident. Under the hypothesis that all human inefficiency is due to noise, stimulus variability must impact human and ideal observers identically: the stimulus-driven variance in the human decision variable (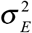 in Eq. 2) will equal the stimulus-driven variance in the ideal observer decision variable (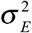 in Eq. 1). Plugging Eq. 1 into Eq. 2 shows that, under the stated hypothesis, human decision variable correlation equals efficiency

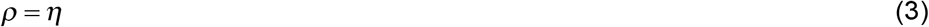

This mathematical relationship has important consequences. It means that the estimate of human efficiency from the speed discrimination experiment (Fig. 4) provides a zero-free parameter prediction of human decision variable correlation in the double pass experiment (Fig. 5). The behavioral data confirm this prediction. Human efficiency in the discrimination experiment quantitatively predicts human response agreement in the double-pass experiment (Fig. 5C; symbols vs. solid curve). The implication of this result is striking. It suggests that naturalistic stimulus variability equally limits human and ideal observers and thus that the source of human inefficiency is due near-exclusively to late noise. Human speed discrimination is therefore optimal except for the impact of late internal noise.

These results generalize across all conditions and human observers. Fig. 6A shows response agreement vs. proportion comparison chosen for the first human observer in each of the five standard speed conditions. (Data from all human observers are shown in Fig. S7.) Fig. 6B summarizes the agreement data across all standard speeds for each human observer. Measured agreement is plotted against the agreement predicted by efficiency. 95% confidence intervals are also shown, which represent prediction uncertainty given the number of double-pass trials in each condition. The decision variable correlations that best account for the response repeatability across all conditions of the first, second, and third human observers are 0.45, 0.43, and 0.18, respectively (see Figs. S7–S8). For the first two observers, stimulus-driven variance and noise variance have approximately same magnitude. For all observers, the data is consistent with the hypothesis that decision variable correlation equals efficiency (solid curves), and is not consistent the null model in which decision variable correlation equals zero (dashed curves). Fig. 6C plots decision variable correlation against efficiency for each human observer. Efficiency tightly predicts decision variable correlation for all three human observers, with zero additional free parameters.

**Figure 6.**
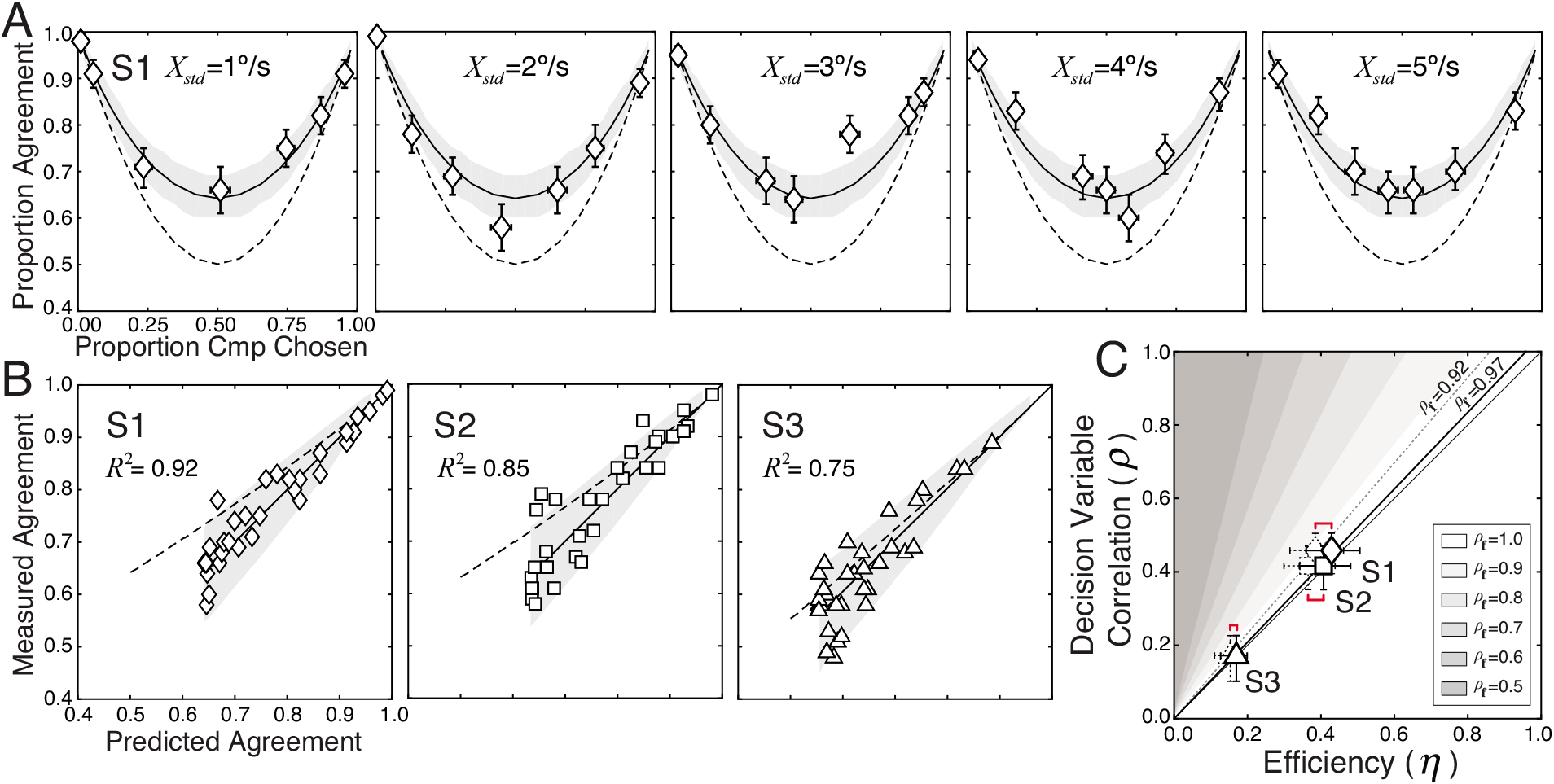
Predicted vs. measured response agreement and decision variable correlation. **A** Proportion response agreement vs. proportion comparison chosen for all five standard speeds (1-5deg/sec), for the first human observer. Human data (symbols) and predictions (curves) are shown using the same conventions as Fig. 5D. **B** Measured vs. predicted response agreement for all conditions and all human observers (symbols). Human agreement equals efficiency-predicted agreement for all three human observers (solid line); shaded regions indicate 95% confidence intervals on the prediction from 1000 Monte Carlo simulations. Efficiency-predicted agreement for the null model, which assumes decision variable correlation is zero, is also shown (dashed curve). **C** Decision variable correlation vs. efficiency for each human observer (symbols). Human efficiency, measured in first pass of the speed discrimination experiment, tightly predicts human decision variable correlation in the double pass experiment with zero free parameters. Error bars represent 95% bootstrapped confidence intervals on human efficiency and on human decision variable correlation. Shaded regions show the expected relationship between efficiency and decision variable correlation if humans use fixed suboptimal computations (i.e. sub-optimal receptive fields). Solid and dashed black lines are the best-fit regression lines, corresponding to receptive field correlations of 0.97 and 0.92, respectively. Red brackets indicate uncertainty about the precise value of efficiency due to uncertainty about the precise amount of early noise (see Fig. 2).

These results indicate that, in two of three observers, naturalistic stimulus variability accounts for nearly half of all behavioral variability. Furthermore, the stimulus set that we used to probe speed discrimination performance almost certainly underestimates the importance of stimulus variability in natural viewing (see Discussion). These considerations suggest that in many tasks, the dominant performance-limiting factors may be external to the observer in natural viewing^19^.

### Suboptimal computations

Human efficiency predicts human decision variable correlation, but to strongly conclude that human inefficiency is due to late noise, it is important to verify that this result is inconsistent with other sources of inefficiency. What is the quantitative impact of fixed suboptimal computations on stimulus-driven variability? To address this question, we analyzed the estimates of a degraded observer that uses suboptimal receptive fields^7,38–40^. If the wrong features are encoded, informative features may be missed, irrelevant features may be processed, and the variance of the stimulus-driven component of the decision variable may be increased relative to the ideal. To create suboptimal receptive fields, we corrupted the optimal receptive fields with fixed samples of Gaussian white noise. Receptive field correlation (i.e. cosine similarity) quantifies the degree of sub-optimality 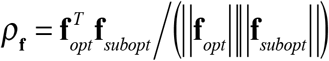 where **f**_*opt*_ and **f**_*subopt*_ are the optimal and suboptimal receptive fields, respectively. We generated degraded observers with different receptive field correlations and examined estimation performance. We found that stimulus-driven variance 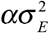 of degraded observer estimates is a scaled version of the ideal stimulus-driven variance. The scale factor 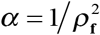 is equal to the squared inverse of receptive field correlation (Fig. S9). Thus, suboptimal receptive fields systematically increase the variance of the stimulus-driven component of the decision variable.

How do fixed suboptimal computations impact the relationship between efficiency and decision variable correlation? If humans are well modeled by a degraded observer with both late noise and suboptimal receptive fields, the total variance of the human estimates is given by 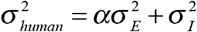. Replacing terms in Eqs. 1 and 2 and performing some simple algebra shows that the relationship between efficiency and decision variable correlation is given by

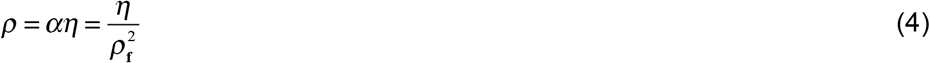

Thus, with sub-optimal computations (i.e. receptive fields) decision variable correlation will be systematically larger than efficiency. This expression has been verified by simulation and a derivation from first principles (Fig. S10; see Supplement). (Note that when receptive field correlation equals 1.0, Eq. 4 reduces to Eq. 3.) We reanalyzed results in the context of Eq. 4, comparing the behavioral data to the predictions of various degraded observer models. For all three observers, decision variable correlation is larger than efficiency by 5%, corresponding to receptive field correlations of 0.97 (Fig. 6C). (Note that these numbers assume an ideal observer with early noise at the upper bound set by the detection experiment (see Fig. 2; Fig. S2). If zero noise is assumed, decision variable correlation exceeds efficiency by 15%, corresponding to receptive field correlation of 0.92.) The vast majority of human inefficiency (minimum 85%) is therefore attributable to late internal noise and not suboptimal feature encoding.

## DISCUSSION

Simple stimuli and/or simple tasks have dominated behavioral neuroscience because of the need for rigor and interpretability in assessing stimulus influences on neural and behavioral responses^41^. The present experiments demonstrate that, with appropriate techniques, the required rigor and interpretability can be obtained with naturalistic stimuli. We have shown that image-computable ideal observers can be fruitfully combined with human behavioral experiments to reveal the factors the limit behavioral performance in mid-level tasks with natural stimuli. In particular, an image-computable ideal observer, constrained by the same factors as the early visual system, predicts the pattern of human speed discrimination performance with naturalistic stimuli^13^. Perhaps more remarkably, human efficiency in the task predicts human decision variable correlation in a double pass experiment without free parameters, a result that holds only if the deterministic computations performed by humans are very nearly optimal.

### Stimulus variability and behavioral variability

A fundamental premise of this paper is that natural stimulus variability limits behavioral performance and drives response repeatability. If this premise is correct, reducing stimulus variability should increase behavioral performance but decrease response repeatability. To test this prediction, we ran a new speed discrimination experiment using drifting random-phase sinewave gratings (Fig. 7; Fig. S11), a stimulus set with less variability than the set of naturalistic stimuli used in the main experiment. As predicted, human speed discrimination thresholds improve (Fig. 7A), responses become less repeatable (Fig. 7B), and decision variable correlation is systematically lower with sinewave stimuli (Fig. 7C). With reduced stimulus variability, internal noise—which is uncorrelated across stimulus repeats—becomes the dominant source of variability limiting human performance.

**Figure 7.**
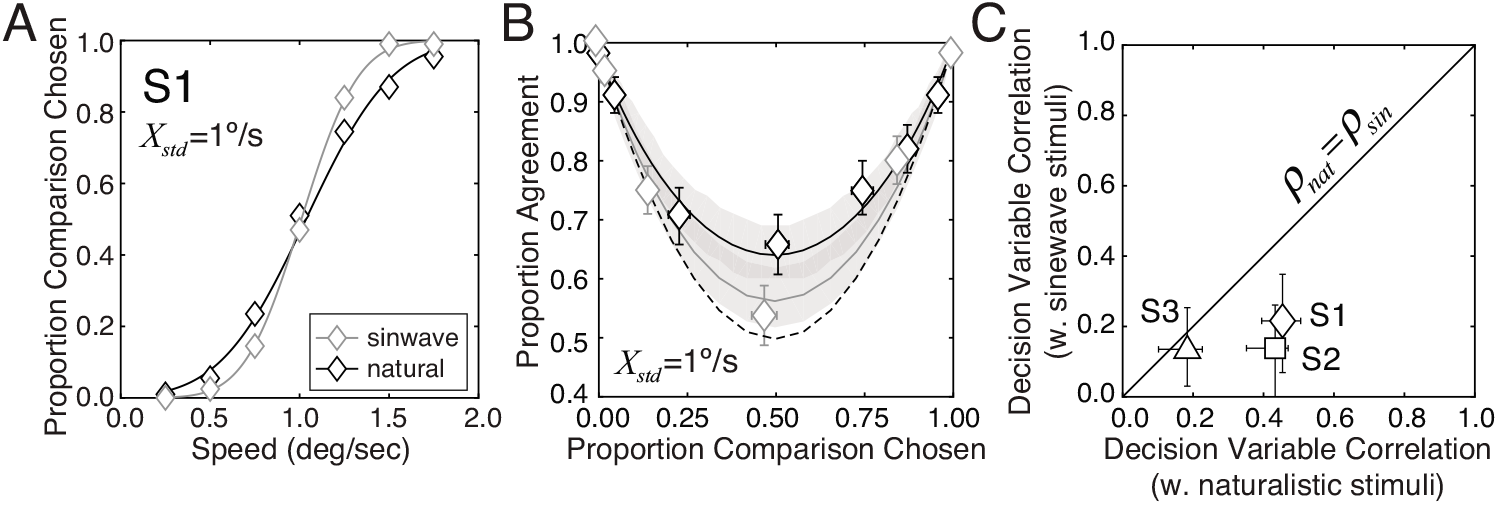
Effects of reducing stimulus variability. **A** Speed discrimination psychometric functions for the first human observer with naturalistic stimuli (black curve) and drifting sinewave stimuli (gray curve) for a 1 deg/sec standard speed. Sinewave stimuli can be discriminated more precisely. **B** Proportion response agreement vs. proportion comparison chosen for naturalistic stimuli (black) and artificial stimuli (grey) for the same human observer. **C** Decision variable correlation with artificial stimuli vs. decision variable correlation with naturalistic stimuli for each human observer (symbols). Error bars represent 95% bootstrapped confidence intervals. Decision variable correlation is consistently lower when artificial stimuli are used. Note that reducing stimulus variability affects the decision variable correlation S3 less than it does observers S1 and S2. This is the expected pattern of results given that S3 had low decision variable correlation with naturalistic stimuli and was thus already dominated by internal noise (see Fig. 6C).

### Limitations and future directions

One limitation of our approach, which is common to most psychophysical approaches, is that it cannot pinpoint the processing stage or brain area at which the limiting source of internal noise arises. Although we model it as occurring at the level of the decision variable, it could also occur at the encoding receptive field responses, the computation of the likelihood, the readout of the posterior into estimates, the placement of the criterion at the decision stage, or some combination of the above. We have ruled out the possibility that the noise limiting speed discrimination is early(Fig. 3; Fig. S2). But we cannot pinpoint where the limiting noise arises. Another limitation is that the approach cannot distinguish between different types of suboptimal computations. We modeled them by degrading each in the set of optimal receptive fields. But an array of computations that make suboptimal use of the available stimulus information could have similar effects. The answers to these questions are best addressed with neurophysiological methods.

There are many possible directions for future work. First, there is a well established tradition of examining how changes in overall contrast impact speed sensitive neurons and speed perception^4,42–47^. All stimuli in the current experiment were fixed to the most common contrast in the natural image movie set. As overall contrast is reduced speed sensitive neurons respond less vigorously, and moving stimuli are perceived to move more slowly^42–44^. It is widely believed that these effects occur because the visual system has internalized a prior for slow speed^43^. In the current manuscript, rather than covering well-trodden ground, we have focused on quantifying how image structure (i.e. the pattern of contrast) impacts speed estimation and discrimination. Thus, our results likely underestimate the impact of stimulus variability on ideal and human performance in natural viewing. The approach advanced in this manuscript can be generalized to examine how changes in overall contrast impact human and ideal performance. The role of stimulus variability has not been examined in this context. A thorough investigation of this issue is an important topic for future work. Finally, experiments should performed with with full space-time (i.e. *xyt*) movies, with stimuli containing looming and discontinuous motion^16,17^, and in a host of new tasks in vision and in other sensory modalities. New databases of natural images and sounds with groundtruth information about distal scenes will significantly aid these efforts^48–50^.

### Sources of performance limits

Efforts to determine the dominant factors that limit performance span research from sensation to cognition. The conclusions that researchers have reached are as diverse as the research areas in which the efforts have been undertaken. Stimulus noise^19^, physiological optics^8^, internal noise^7,20–22^, suboptimal computations^39,51,52^, trial-sequential dependences^53^, and various cognitive factors^54^ have all been implicated as the dominant factors that limit performance. What accounts for the diversity of these conclusions? We cannot provide a definitive answer, but we speculate that the relative importance of these factors is likely to depend on several factors.

Evolution has pushed sensory-perceptual systems towards the optimal solutions for tasks that are critical for survival and reproduction. Humans are more likely to be assessed as optimal when visual systems are probed with stimuli that they evolved to process in tasks that they evolved to perform. In target detection tasks, for example, humans become progressively more efficient as stimuli become more natural^8,55,56^. Conversely, when stimuli and tasks bear little relation to those that drove the evolution of the system, the computations are less likely to be optimal. This issue suggests that a new scientific framework that takes these factors into account may help to reconcile these disparate findings.

### Image-computable ideal observers

Ideal observer analysis has a long history in vision science and systems neuroscience. In conjunction with behavioral experiments, image-computable ideal observers have shown that human light sensitivity is as sensitive as allowed by the laws of physics^19^, that the shape of the human contrast sensitivity function is dictated by the optics of the human eye^8^, and that the pattern of human performance in a wide variety of basic psychophysical tasks can be predicted from first principles^9^.

To develop an image-computable ideal observer, it is critical to have a characterization of the task-relevant stimulus statistics. Obtaining such a characterization has been out of reach for all but the simplest tasks with the simplest stimuli. The vision and systems neuroscience communities have traditionally focused on understanding how simple forms of stimulus variability (e.g. Poisson or Gaussian white noise) impact performance^7,8,19,20,57^. The impact of natural stimulus variability—the variation in light patterns associated with different natural scenes sharing the same latent variable values—has only recently begun to receive significant attention^6,10–13,56,58–62^.

Many impactful ideal observer models developed in recent years are not image-computable^43,63–67^. The weakness of these models is that they do not explicitly specify the stimulus encoding process, and therefore make assumptions about the information that stimuli provide about the task relevant variable (e.g. the likelihood function in the Bayesian framework). In other words, these models cannot predict directly from stimuli how stimulus variability will impact behavioral variability. Image-computable models are thus necessary to achieve the goal of understanding how vision works with real-world stimuli. The current work represents an important step in that direction.

### Impact on neuroscience

Behavioral and neural responses both vary from trial to trial even when the value of the latent (e.g. speed) is held constant. In many classic neurophysiological experiments, stimulus variability is eliminated by design, and experimental distinctions are not made between the latent variable of interest (e.g. orientation) and the stimulus (e.g. an oriented Gabor) used to probe neural response. Such experiments are well suited for quantifying how different internal factors impact neural variability. Indeed, it has recently been shown that, under these conditions, neural variability can be partitioned into two internal factors: a Poisson point-process and system-wide gain fluctuations^68^. This approach provides an elegant account of a widely observed phenomenon (‘super-Poisson variability^69–71^) that had previously resisted rigorous explanation. However, the designs of these classic experiments are unsuitable for estimating the impact of stimulus variability on neural response.

In the real world, behavioral variability is jointly driven by external and internal factors. Our results show that both factors place similar limits on performance. A full account of neural encoding and decoding must include a treatment of all significant sources of response variability. Partitioning the impact of realistic forms of stimulus variability from internal sources of neural variability will be an important next step for the field.

### Author Contributions

BMC and JB conceived the project, collected and analyzed data, performed computational analyses, and wrote and edited the paper.

## Acknowledgments

This work was supported by startup funds to JB from the University of Pennsylvania, and by NIH grant R01-EY028571 to JB from the National Eye Institute and the Office of Behavioral and Social Sciences Research.

## Data availability

The data supporting the findings of this study are available on reasonable request.

## Code availability

The code used to analyze the data of this study is available on reasonable request.

## Methods

### Human observers

Three observers participated in the experiment. All had normal or corrected-to-normal acuity.

### Equipment

Stimuli were presented on a ViewSonic G220fb 40.2cm × 30.3cm cathode ray tube monitor with 1280×1024pixel resolution, and a refresh rate of 60 Hz. At the at the 92.5 cm viewing distance, the monitor subtended a field of view of 24.5° × 18.6° of visual angle. The display was linearized over 8 bits of grey level. The maximum luminance was 74 cd/m^2^. The mean background grey level was set to 37 cd/m^2^. The observer’s head was stabilized with a chin-and-forehead rest.

### Stimuli: Detection experiment

Target stimuli in the detection experiment consisted of static, vertically-oriented Gabor targets in cosine-phase (3cpd and 4.5cpd) with 1.5 octave bandwidths embedded in vertically-oriented (1D) dynamic Gaussian noise. Targets subtended 1 deg of visual angle for a duration of 250ms (15 frames at 60htz). Stimuli were windowed with a raised-cosine window in space and a flattop-raised-cosine window in time, exactly the same as the image movies in the speed discrimination experiment. The RMS contrast of the target and the noise were varied independently according to the experimental design. To minimize target uncertainty, the target was presented to the subject, without noise every 10 trials.

For the detection experiment, a bit-depth of greater than 8 bits is required to accurately measure contrast detection thresholds. We achieved a bit-depth of more than 10 bits using the LOBES video switcher^72^. The video switcher combines the blue channel and attenuated red channel outputs in the graphics card. Picking the right combination of blue and red channel outputs generates a precise gray-scale luminance signal.

### Procedure: Detection experiment

Stimuli in the target detection experiment were presented using a two-interval forced choice (2IFC) procedure. On each trial, one interval contained a target plus noise, and the other interval contained noise only. The task was to select the interval containing the target. Feedback was provided. Psychometric functions were measured for each of four different noise contrasts (0.00, 0.05, 0.10, 0.20) using the method of constant stimuli, with five different target contrasts per condition. Each observer completed 3200 trials in this experiment (4 noise levels × 5 target contrasts per noise level × 80 trials per target × 2 target frequencies). Each block contained 50 trials. To minimize observer uncertainty trials were blocked by stimulus and noise contrast. The target stimulus was also presented at the beginning of each block, and then again every 10 trials, throughout the experiment.

### Stimuli: Speed discrimination experiment

Natural image movies were created by texture-mapping randomly selected patches of calibrated natural image onto planar surfaces, and then moving the surfaces behind a stationary 1.0° aperture. The movies were then restricted to one dimension of space by vertically averaging each frame of the movie^13^. Each movie subtended 1 deg of visual angle. Movie duration was 250ms (15 frames at 60htz). All stimuli were windowed with a raised-cosine window in space and a flattop-raised-cosine window in time. The transition regions at the beginning and end of the time window each consisted of four frames; the flattop of the window in time consisted of seven frames. Contrast was computed under the space-time window. To prevent aliasing, stimuli were low-pass filtered in space and time before presentation (Gaussian filter in frequency domain with *σ_space_* =4cpd, *σ_time_* =30htz). No aliasing was visible.

All stimuli were set to have the same mean luminance as the background and had a fixed root-mean-squared (RMS) contrast of 0.14 (equivalent to 0.20 Michelson contrast for sinewave stimuli), the modal contrast of the stimulus ensemble. The RMS contrast is given by

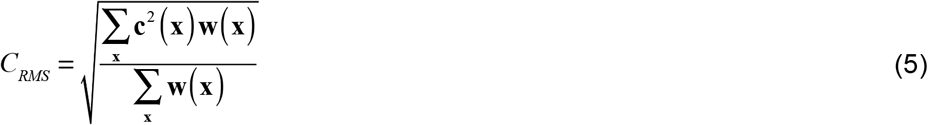

where **c**(**x**) is a Weber contrast image movie, **w**(**x**) is the window, and **x** = {*x,y,t*} is a vector of space-time positions. Stimuli were contrast fixed because contrast is known to effect speed percepts and our focus was on how differences in Weber contrast patterns between stimuli impact performance rather than on differences in overall contrast impact performance, which have already been intensively studied^42,43^.

### Procedure: Speed discrimination experiment

For the speed discrimination task, data was collected using a 2IFC procedure. On each trial, a standard and a comparison image movie were presented in pseudo-random order (see below). The task was to choose the interval with the movie having the faster speed. Human observers indicated their choice via a key press. The key press also initiated the next trial. Feedback was given. A high tone indicated a correct response; a low tone indicated an incorrect response. Experimental sessions were blocked by absolute standard speed. In the same block, for example, data was collected at the −5 and +5 deg/sec standard speeds. Movies always drifted in the same direction within a trial, but directions were mixed within a block. An equal number of left- and right-drifting movies were presented in the same block to reduce the potential effects of adaptation.

In each pass of the experiment (see below), psychometric data were measured for each of 10 standard speeds (+5, +4, +3, +2, +1) using the method of constant stimuli, with seven comparison speeds per function. For each standard, each of the seven comparison speeds was presented 50 times. Thus, on each pass, each observer completed 3,500 trials (2 directions × 5 standard speeds × 7 comparison speeds × 50 trials).

The exact same naturalistic movie was never presented twice within a pass of the experiment. Rather, movies were randomly sampled without replacement from a test set of 1,000 naturalistic movies at each speed. For each standard speed, 350 ‘standard speed movies’ were randomly selected. Similarly, for each of the seven comparison speeds corresponding to that standard, 50 ‘comparison speed movies’ were randomly selected. Standard and comparison speed movies were then randomly paired together. This stimulus selection procedure was used to ensure that the stimuli used in the psychophysical experiment had approximately the same statistical variation as the stimuli that were used to train and test the ideal observer model. Assuming the stimulus sets are representative and sufficiently large, the stimuli presented in the experiment are likely to be representative of natural signals.

### Ideal observer for speed estimation

As signals proceed through the visual system, neural states become more selective for properties of the environment, and more invariant to irrelevant features of the retinal images. The ideal observer for speed estimation computes the Bayes’ optimal speed estimate from the posterior probability distribution over speed *p*(*X*|**R**) given the responses **R** of a small population of space-time receptive fields to a stimulus^13^. Given the constraints imposed by natural stimulus variability, measurement noise, and the early visual system, the space-time receptive fields and the subsequent computations for decoding the speed must be optimal in order for the estimates to be optimal (Fig. S3A). The most useful stimulus features and the computations that optimally pool them are jointly dictated by the task and the properties of natural stimuli. The receptive fields that encode the most optimal stimulus features for the task are determined via a recently developed technique called Accuracy Maximization Analysis^18,29,30^ (AMA). The optimal computations for pooling the responses of the receptive fields are specified by how the receptive field responses are distributed (Fig. S4B). The conditional receptive field responses *p*(**R**|*X_k_*) = *gauss*(**R;0**,Σ_*k*_) are jointly Gaussian and mean zero^13,18^ after response normalization. For any observed response **R**, the computations that specify the likelihood *L*(*X_u_*;**R**) = *p*(**R**|*X_u_*) that an observed response was elicited by a stimulus moving with speed *X_u_* is obtained by evaluating the response in the response distribution corresponding to that speed. The responses must therefore be pooled in a weighted quadratic sum, with weights that are given by simple functions of the covariance matrices^13^. A neuron that performs these quadratic computations outputs a response 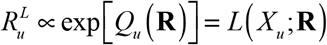 that is proportional to the likelihood that a stimulus moving at speed *X_u_* elicited the response **R**. Following response normalization^26–28^, the likelihood neurons instantiate an energy-model-like hierarchical LNLN (linear, non-linear, etc.) cascade^2,18^. Thus, the computations that yield likelihood neurons can be thought of as a computational recipe, grounded in natural image and scene statistics, for how to optimally construct selective invariant speed-tuned neurons (Fig. S3D).

To obtain the posterior probability of each speed, the likelihood must be weighted by the prior *p*(*X_u_*) and normalized by the weighted sum of likelihoods Σ_*v*_*L*(*X_v_*;**R**)*p*(*X_v_*). Finally, the optimal estimate must be ‘read out’ from the posterior probability distribution. In the case of the 0,1 cost function (i.e. L0 norm) the optimal estimate 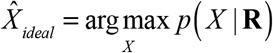 is the posterior max (Fig. S3C). If the prior probability distribution is flat, the optimal estimate is the latent variable value that corresponds to the maximum of the likelihood function (i.e. the max of the likelihood neuron population response; Fig. 3EF).

### Ideal, degraded, and human decision variables

The ideal decision variable for the task of speed discrimination is obtained by the subtracting optimal the speed estimates corresponding to the comparison and standard stimuli

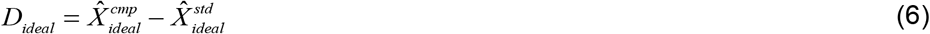

If the decision variable is greater than zero, the ideal observer responds that the comparison stimulus was faster. If the decision variable is less than zero, the ideal observer responds that the comparison stimulus was slower. Degraded observer decision variables are similarly obtained, except that the degraded observer estimates are obtained by reading out the responses of suboptimal receptive fields (Fig. S10).

The human decision variable is a noisy version of the ideal decision variable, under the hypothesis that human inefficiency is due only to internal noise. Specifically,

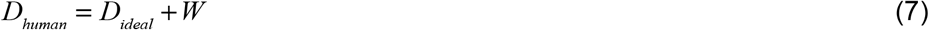

where 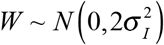 is a sample of zero mean Gaussian noise, which corresponds to adding noise with variance 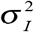 to the comparison and standard stimulus speed estimates.

### Double pass experiment

A double pass experiment requires that each observer performs all (or a subset) of the unique trials in an experiment twice. In our experiment, each trial was uniquely identified by its standard and comparison movies. An observer completed the first pass by completing each unique trial once over 20 blocks consisting of 175 trials each. The standard speed was always constant within a block. Blocks were counterbalanced. The observer completed the second pass by completing each unique trial again over another 10 blocks. Before collecting data in the main experiment, each human observer completed multiple practice sessions to ensure that perceptual learning had stabilized. Analysis of the practice data showed no significant learning effects. Stimuli presented in practice sessions were not presented in the main experiment.

### Estimating decision variable correlation

Human decision variable correlation is estimated via maximum likelihood from the pattern of human response agreement in the double-pass experiment. The log-likelihood of the double-pass response data is given by

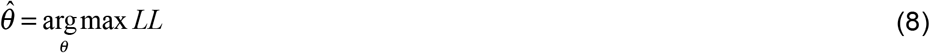

where ***θ*** is a vector of model parameters describing decision variable distribution and observer criteria across both passes of the double pass experiment. The log-likelihood of the double-pass response data is given by

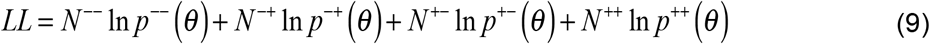

where *N*^−−^, *N*^−+^, *N*^+−^, *N*^++^ are the number of sampled decision variables (i.e. responses) in the lower left, upper left, lower right, and upper right quadrants of the response space (see Figs. 4B, S6B-D). The likelihoods of observing those samples are given by

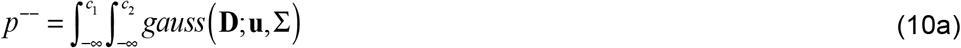

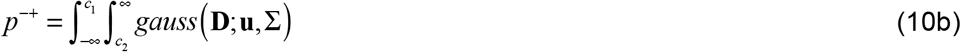

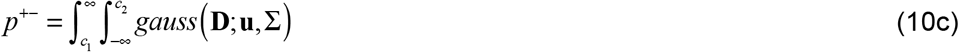

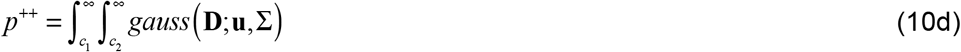

where **D** is the joint decision variable across passes with mean **u** and covariance Σ and *c*_1_ and *c*_2_ are the observer criteria on passes one and two. The mean decision variable values are set equal to the speed difference *μ*_1_=*μ*_2_ = *X_cmp_* − *X_std_* between the standard and comparison stimuli in each condition.

In practice, and without loss of generality, we estimate the decision variable correlation using normalized decision variables **Z**. The parameter vector for maximizing the likelihood of the normalized decision variables is 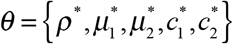 where * indicates that the parameter is associated with the normalized variable. The integrals in Eq. 10a–d can be equivalently expressed with limits of integration *c*\* = *c/σ_human_* and integrand *gauss*(**Z;Mu,MΣM^*T*^**) with normalized mean and normalized covariance

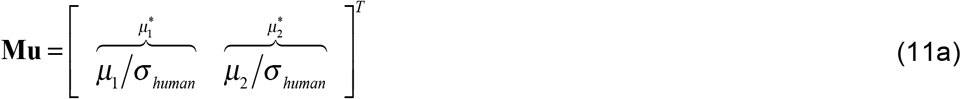

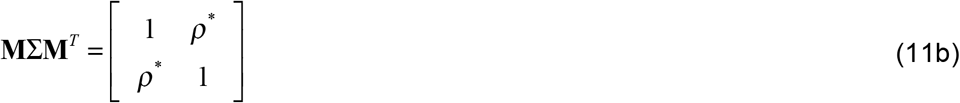

where the normalizing matrix is 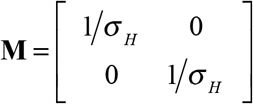. Normalizing the variables has the practical advantage that it converts the covariance matrix to a correlation matrix, so that it can be fully characterized with a single parameter: decision variable correlation. It also sets the normalized means equal to *d*′. We fix the normalized means 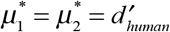 to the human sensitivity measured in the discrimination experiment (c.f. Fig. 4). We also fix the normalized criteria to 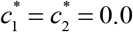, which is justified by the data (Fig. S4). These choices reduce the number of parameters to be estimated from five to one.

### Efficiency and early noise

The approximate equality in Eq. 1 is useful for developing intuitions about the relationship between efficiency and decision variable correlation. But the approximate equality in Eq. 1 assumes that the impact of early noise on the ideal observer decision variable is negligible. The exact expression for efficiency is given by

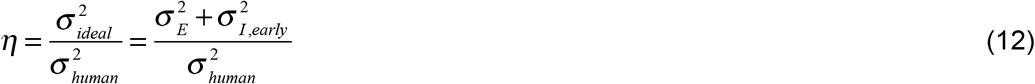

where 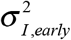, is the variance in the decision variable due to early noise. The variance in the decision variable due to early noise is distinct from early noise itself, which is defined in the domain of the image pixels instead of the decision variable. This is analogous to how the stimulus-driven variance 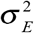 in the decision variable is distinct from stimulus variability. Stimulus variability, like early noise, is defined in domain of the image pixels and is non-zero in any set of non-identical stimuli having the same value of the latent variable. We computed efficiency using the exact expression in Eq. 12 instead of the approximate equality in Eq. 1 and found that, for our estimated values of early noise, both expressions produce similar estimates of efficiency (Fig. S2).

#### Supplement

**Figure S1.**
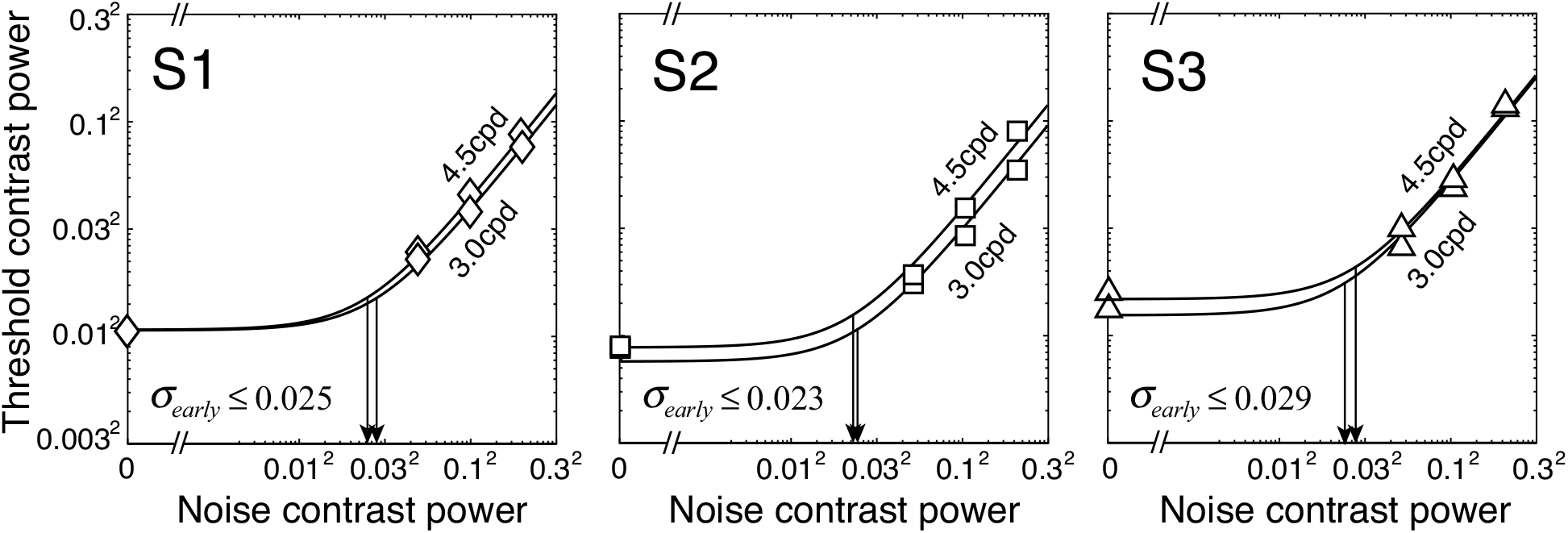
Threshold contrast power (i.e. equivalent input noise) for both targets and all three human observers observers. Equivalent input noise, expressed as percent contrast in the retinal image, is 2.5%, 2.3%, and 2.9% for each observer, respectively. The knee of each function (arrow) indicates the contrast power of the pixel noise required to double the threshold associated with zero pixel noise. Thus, the equivalent input noise bounds the amount of early noise in the system.

#### Estimating early noise

In target detection tasks, in which a known target must be detected in noise, the square of the detection threshold increases linearly with the stimulus noise power^7^. This fact can be leveraged to estimate the internal noise that limits detection performance. Stimulus noise (i.e. pixel noise) is under experimental control. Internal noise is not. Both noise types influence target detection thresholds. Target contrast power at threshold is a function of stimulus noise 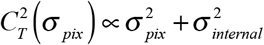 and is proportional to the sum of pixel and internal noise variances. The constant of proportionality depends on the target. Strong inferences can be made about the amount of internal noise from the psychophysical data, by assuming that internal noise is fixed. When pixel and internal noise have equal variance, for example, the squared detection threshold will be twice what it is when pixel noise is zero: 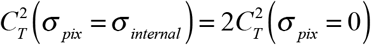. The internal noise limiting performance in a detection experiment can therefore be estimated from the pattern of detection thresholds. The noise that limits detection performance could be early (e.g. at the level of the retinal image), late (e.g. at the level of the decision variable), or some combination of both. The detection thresholds therefore cannot be used to determine the exact amount of early noise. Rather, the detection thresholds place an upper bound on the amount of early noise in each human observer.

**Figure S2.**
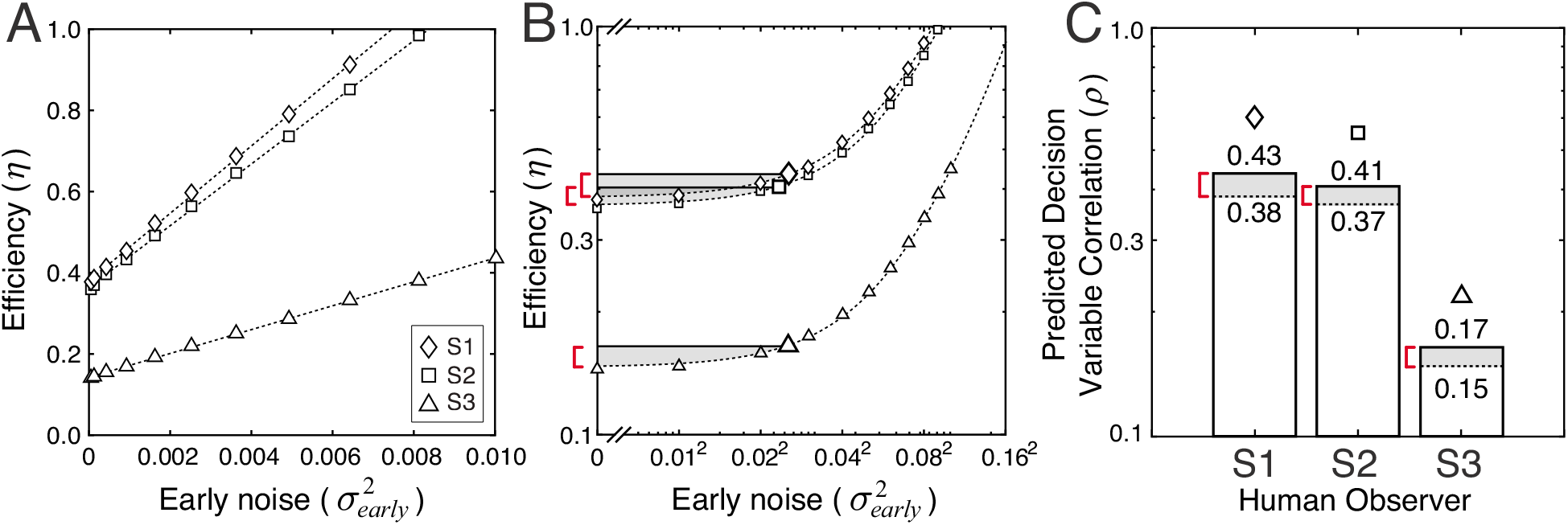
The impact of early noise on ideal observer performance and human efficiency. **A** Efficiency in speed discrimination for each human observer (symbols) as a function of the amount of early noise modeled in the ideal observer. If early noise is negligible, efficiency is given by 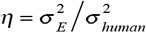. If early noise is non-negligible, efficiency is given by 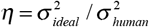 the ratio of the variances of the ideal and human decision variables. The estimate of human efficiency increases linearly with the amount of early noise because increasing the variance of early noise increases the numerator in the expression for efficiency. **B** Same data as A, but on log-log axes. Given the upper bound established by the target detection experiment, the impact of early noise on estimates of human efficiency cannot be large. Throughout the main text, we used an estimate of efficiency that assumes early is at this upper bound (large symbols & solid horizontal lines). If one assumes no early noise (which is logically possible but not biologically plausible), the estimate of efficiency is given by the y-intercept of each curve. The red brackets and shaded areas represent all possible efficiencies consistent with these upper and lower bounds. The true value of efficiency is most likely somewhere in between these maximum and minimum values. **C** Predicted decision variable correlation for each human observer given the uncertainty about human efficiency. The maximum (solid line) and minimum (dashed line) predicted decision variable correlations correspond to ideal observers having the maximum and minimum amount of early noise. The predicted decision variable correlations differ by ~10% at maximum.

#### Early noise and human efficiency

Human efficiency is determined by comparing human and ideal observer sensitivities (i.e. variances; Eq. 1). Human variability is determined by stimulus-driven variability, early noise, and all other types of noise. Ideal variability is determined only by stimulus-driven variability and early noise. If variance of the ideal decision variable is misestimated, the estimate of human efficiency may be inaccurate (Eq. 2). The ideal observer is constrained by the same factors that are known to constrain the human visual system. If the early noise built into the ideal observer is larger than it should be (i.e. more than in the human), ideal variance will be larger than it should be and human efficiency will be over-estimated. If the early noise in the ideal observer is lower than it should be (i.e. less than in the human), ideal variance will be smaller than it should be and human efficiency will be underestimated.

To determine how the amount of early noise impacts the estimate of efficiency, and thus its empirical relationship with decision variable correlation, we systematically varied the amount of early noise in the ideal observer. The amount of early noise ranged from zero, to the upper bound on early noise set by the target detection experiment, to values that exceeded the bound (Fig. S2AB). For each amount of early noise, we recomputed efficiency using the exact equality in Eq. 1 (also see Fig. 12) instead of the approximate equality in Eq. 1 used throughout the manuscript. The estimates of human efficiency with early noise set equal to zero and early noise set equal to the upper bound established by the detection experiment are within ~10% for each human observer (Fig. S2C). Thus, the true value of human efficiency is no more than 10% different than the values reported in the main text.

**Figure S3.**
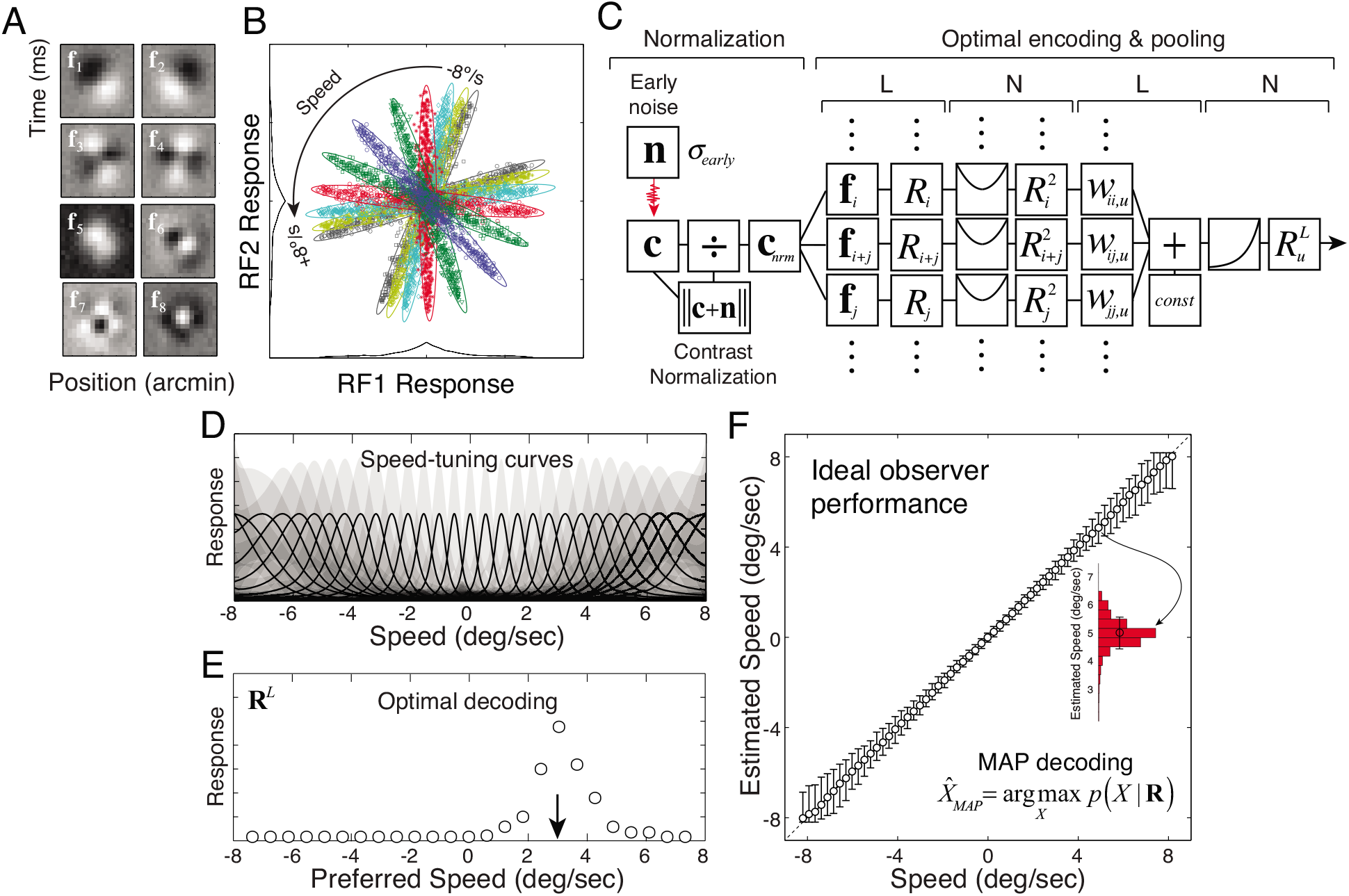
Ideal observer receptive fields, response distributions, estimates, and pooling. **A** Optimal space-time receptive fields (RFs) for speed estimation given the natural stimulus set and the constraints of the early visual system. Eight RFs capture essentially all of the task-relevant information. **B** Receptive field response distributions, conditioned on the speed of the image movie (colors). The joint response of the filters to each stimulus is given by **R** = **f**^*T*^(**c + n**)/||**c + n**|| where **f** is the set of filters, **c** is the contrast stimulus, and **n** is a sample of early noise. Each symbol represents the joint response to an individual movie. Each response distribution is well described by a zero-mean Gaussian *p*(**R**|*X_k_*) = *gauss*(**R**;0,Σ_*k*_). Response distributions for other receptive field pairs are similarly distributed. The response variability within each color is due to natural stimulus variability; that is, it is the stimulus variability in the feature space defined by the optimal RFs. The covariance Σ_*u*_ of the RF responses to movies having speed *X_u_* dictate the optimal pooling weights **w**_*u*_ (see below). **C** Hypothetical neuron implementing optimal encoding and pooling. Each noisy, contrast-normalized stimulus is processed by the optimal RFs. The responses of these RFs are pooled in a weighted quadratic sum. The sum is then pushed through an exponential output nonlinearity to obtain the likelihood *R^L^*(*X_u_*) = *L*(*X_u_*;**R**) that the observed receptive field responses **R** were elicited by speed *X_u_*, the preferred speed of the neuron. The response of this hypothetical neuron represents the likelihood that a given stimulus had its preferred speed. The optimal pooling rules thus represent a LNLN (linear, non-linear, etc.) cascade. **D** Speed tuning curves of hypothetical neurons implementing optimal encoding and pooling, whose responses represent the likelihood of each speed given a stimulus. The speed-tuning curve 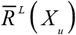 is the average likelihood across stimuli at each of many different speeds. Shaded regions indicate +1SD confidence intervals on response. Response variability is due primarily to natural stimulus variability. The optimal computations convert the tuning curves of the space-time RFs (A), which individually provide poor information about speed, into the speed tuning curves of the likelihood neurons, which individually provide excellent information about speed. **E** An arbitrary stimulus creates a population response **R**^*L*^ over hypothetical speed-tuned neurons. Optimal decoding yields the optimal estimate. **F** Ideal observer estimates. The optimal estimate is read out from the population of hypothetical speed-tuned neurons in E, and is equivalent to reading out the posterior probability distribution *p*(*X*|**R**) over speed. Estimate variability (histogram) is dominated by stimulus-driven variability (***σ***_*E*_; see Figs. 1A, 4B).

**Figure S4.**
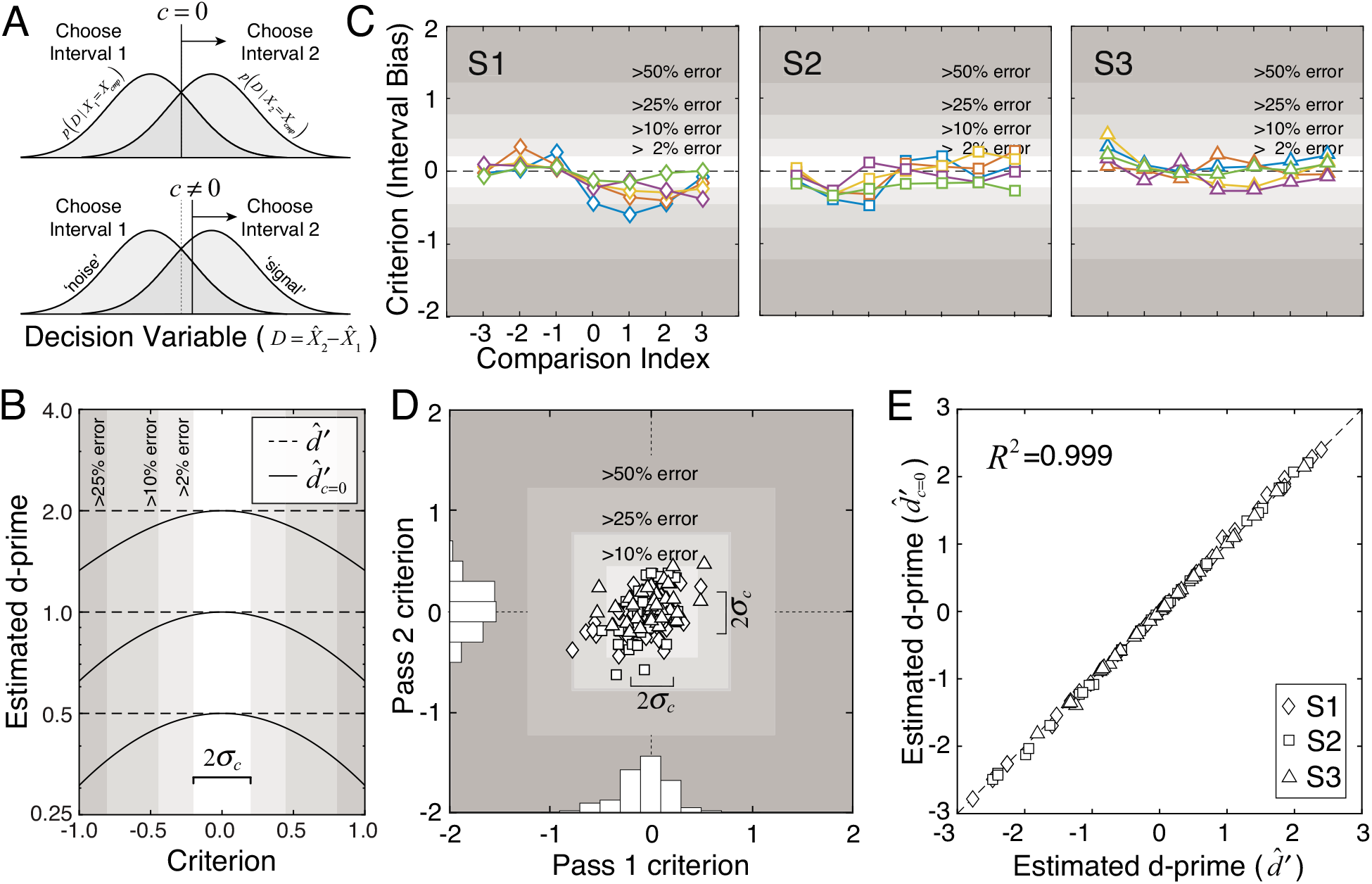
Human criteria and the estimation of d-prime in the speed discrimination experiment. **A** Decision variable distributions with optimal and non-optimal criteria. The decision variable 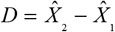 is given by the differences between the speed estimates on the two intervals of each trial. Assuming the comparison speed is greater than the standard speed *X_cmp_* > *X_std_*, the ‘signal’ and ‘noise’ distributions-- *p*(*D*|*X*_2_ = *X_cmp_*) and *p*(*D*|*X*_1_ = *X_cmp_*)--correspond to when the comparison is in the second and first interval, respectively. When the decision variable exceeds the value of the criterion, the observer chooses interval two as faster. If the criterion is optimally placed (i.e. *c* =0), the proportion of hits and correct rejections *p*(*HT*) = *p* (*CR*) equal one another. If the criterion is non-optimally placed (i.e. *c* ≠ 0.0), the proportion of hits and correct rejections do not equal one another *p*(*HT*) ≠ *p*(*CR*). **B** Estimating d-prime with and without the assumption that the criterion equals zero. The expression 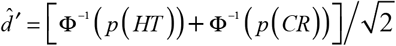 provides an estimate of d-prime that makes no assumption about the location of the criterion (dashed lines). The expression 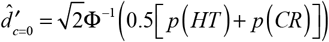 provides an estimate of d-prime assuming that the criterion equals zero (solid curves). For simplicity, the latter expression is used throughout the main text. If the criterion is non-zero, but 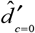 is used, d-prime is systematically underestimated. However, for small criterion shifts, estimation errors are small (shaded regions). **C** Criteria of human observers as a function of the comparison index for each standard speed (colors). Each comparison index corresponds to a comparison speed, three slower and three faster than the standard. Comparison indices are plotted rather than comparison speeds because comparison speeds and how much they differ from the standard speed change with the standard speed. Thirty-five criteria are computed for each observer (5 standard × 7 comparison speeds), averaged across both passes in the experiment. Shaded regions show criteria values for which the d-prime estimation error is greater than 2%, 10%, 25%, and 50%, respectively. **D** Observer criteria on each pass of the experiment for the three observers (symbols). Histograms show the distribution of criteria in each pass. Twice the standard deviation of the criteria *σ_c_* in each pass of the experiment equals the width of the 2% error region, and is half the width of the 10% error region. **E** D-prime computed two ways. D-prime estimated with the assumption that observer criteria equaled zero vs. d-prime estimated without that assumption. The differences are negligible. The approach taken in the main text is therefore justified.

**Figure S5.**
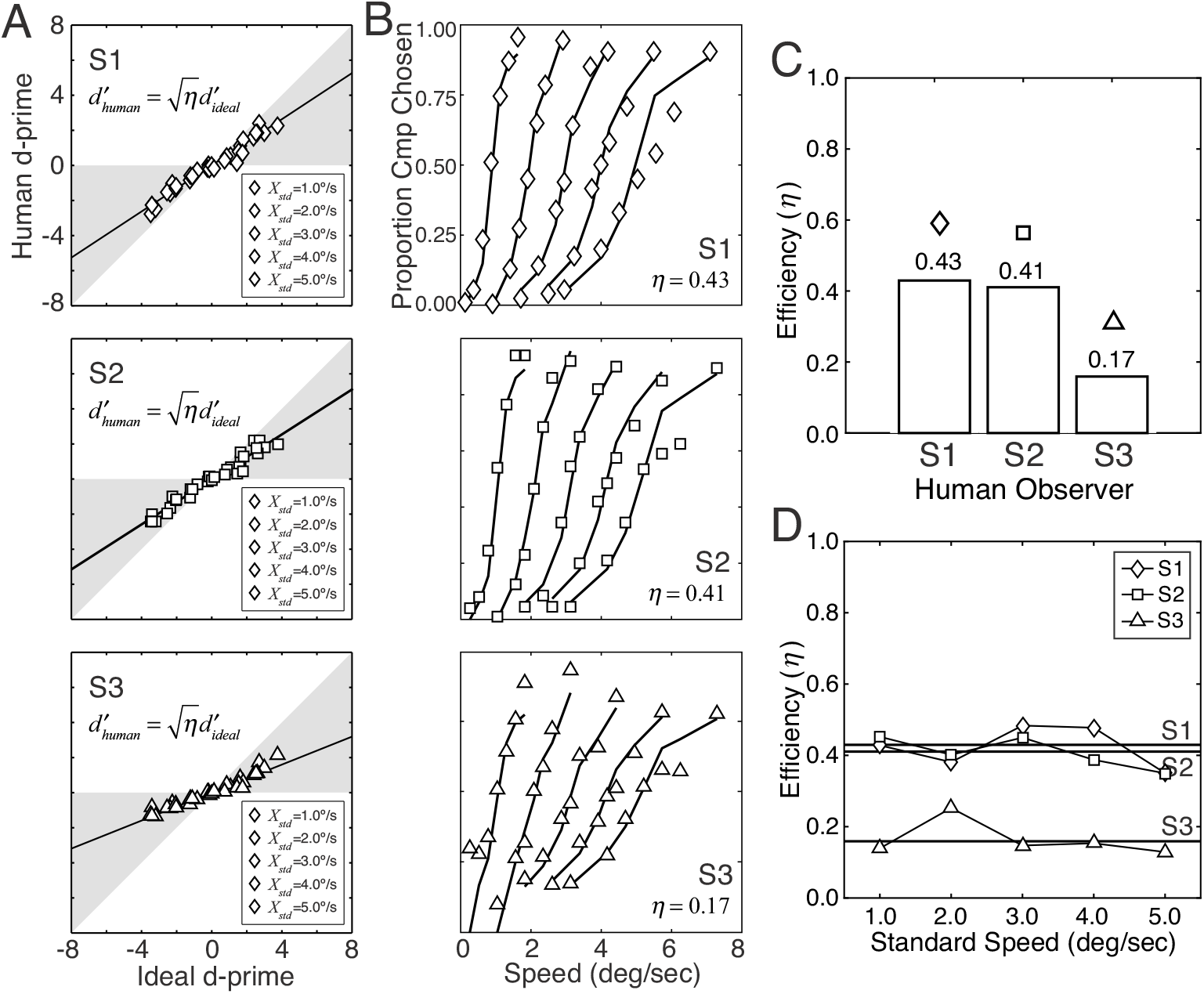
Human sensitivity, psychometric functions, and efficiency vs. the ideal observer. **A** Human vs. ideal observer sensitivity (d-prime) for all three human observers. Human and ideal sensitivities are linearly related by the square root of efficiency. **B** Human psychometric functions (symbols) and degraded ideal observer fits (curves). The degraded ideal observer is fit to each human observer with one free parameter: efficiency. **C** Human efficiencies are 0.43, 0.41, and 0.17. **D** Human efficiency as a function of speed. Efficiency for each human observer is approximately constant with speed.

**Figure S6.**
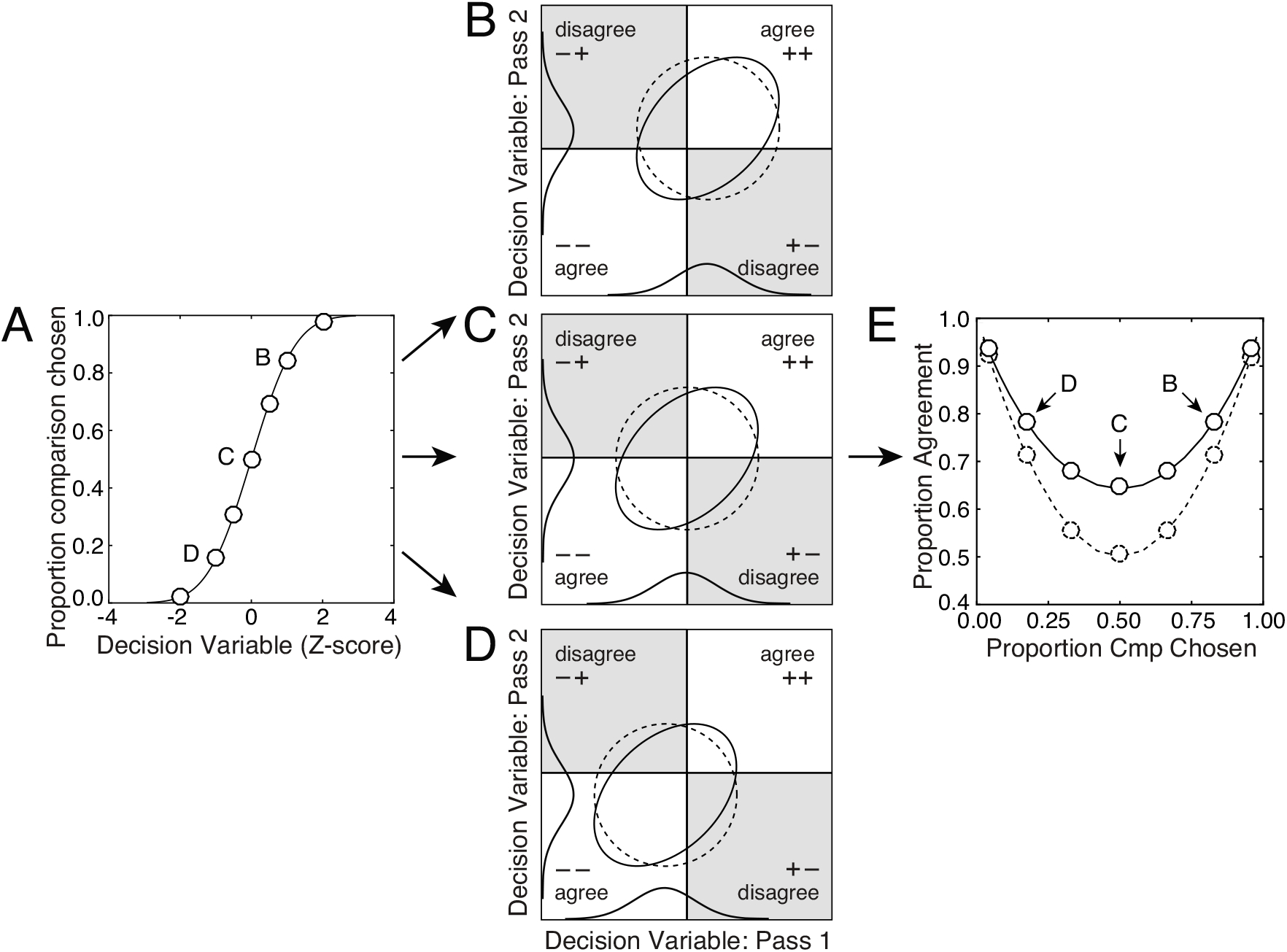
The psychometric function, decision variable correlation, and response agreement. **A** The psychometric function plots proportion comparison chosen vs. the decision variable. **B** Decision variable correlation when proportion comparison chosen is greater than 50%, **C** equals 50%, **D** and is less than 50%. Ellipses representing distributions with decision variable correlations of 0.5 (solid ellipse) and 0.0 (dashed circle) are shown. Response agreements occur when the decision variable is correct on both passes (upper right quadrant) or incorrect on both passes (lower left quadrant). Thus, increased correlation is associated with increased agreement. Note that we depict a single decision variable distribution rather than the two that are standard in signal detection theory. This choice is justified if observer criterion is optimally placed; that is, when the criterion is optimally placed, proportion hits equals proportion correct rejections. In our dataset, non-optimal criterion placement is negligible (see Fig. S4). **E** Proportion response agreement vs. proportion comparison chosen against predictions for decision variable correlations of 0.5 (solid curve) and 0.0 (dashed curve).

**Figure S7.**
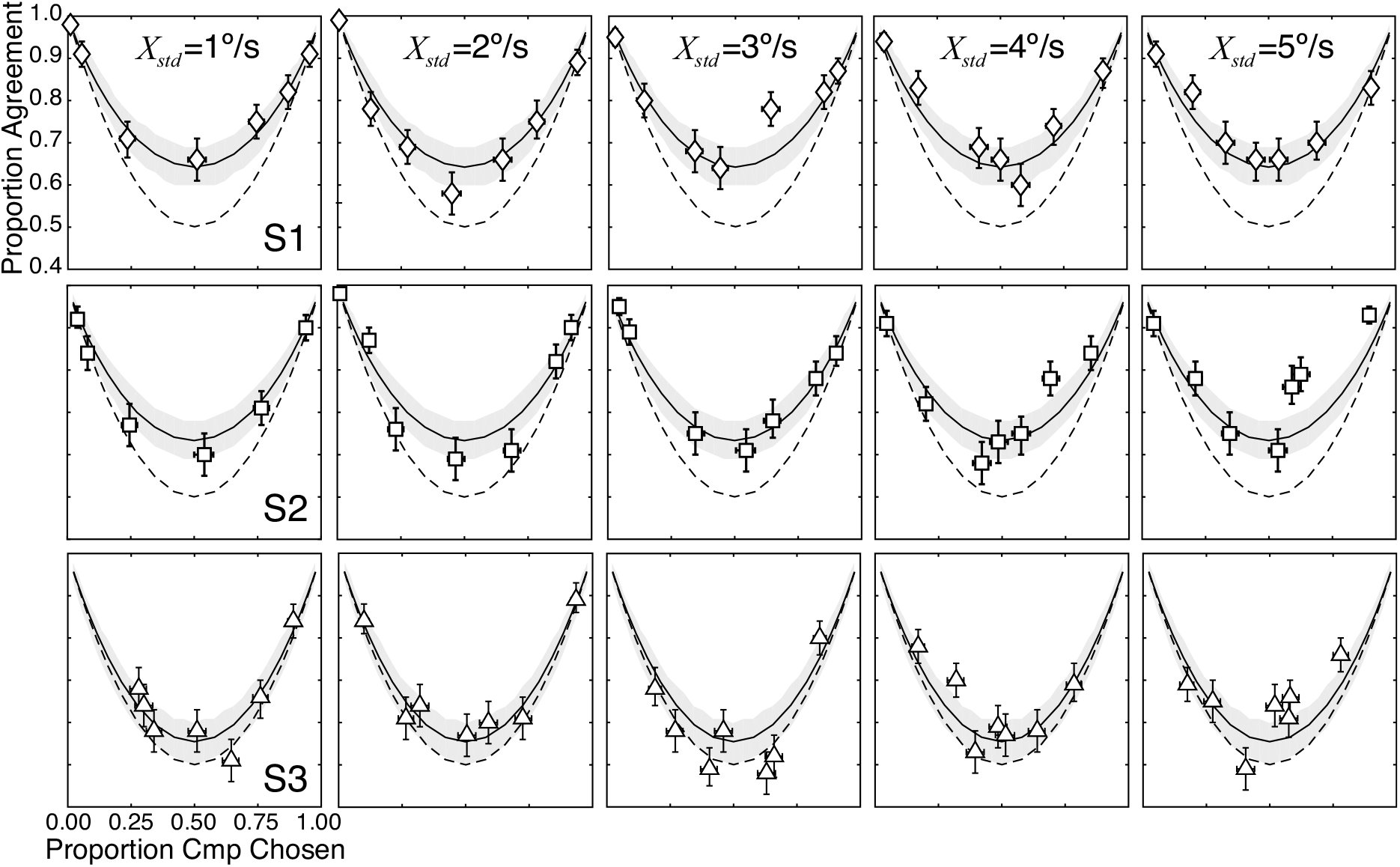
Proportion response agreement vs. proportion comparison chosen in all conditions, for each human observer. Top, middle, and bottom rows show proportion response agreement vs. proportion comparison chosen for human observers S1, S2, and S3, respectively for each of five standard speeds (1-5 deg/sec; columns). Error bars indicate 68% bootstrapped confidence intervals on the agreement data. Shaded regions represent prediction uncertainty (68% confidence intervals) from 10000 Monte Carlo simulations given the uncertainty in the estimate of efficiency.

**Figure S8.**
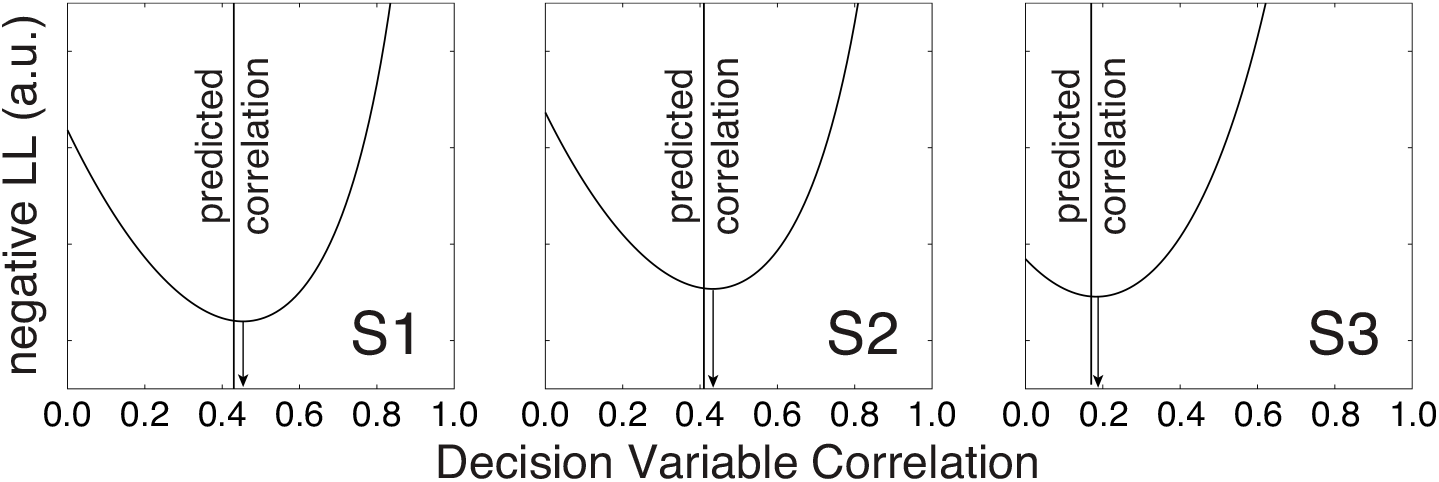
Negative log-likelihood as a function of fitted decision variable correlation for each human observer. The vertical line represents the efficiency-predicted value of decision variable correlation (zero free parameters), under the assumption that all inefficiency is due to internal noise.

**Figure S9.**
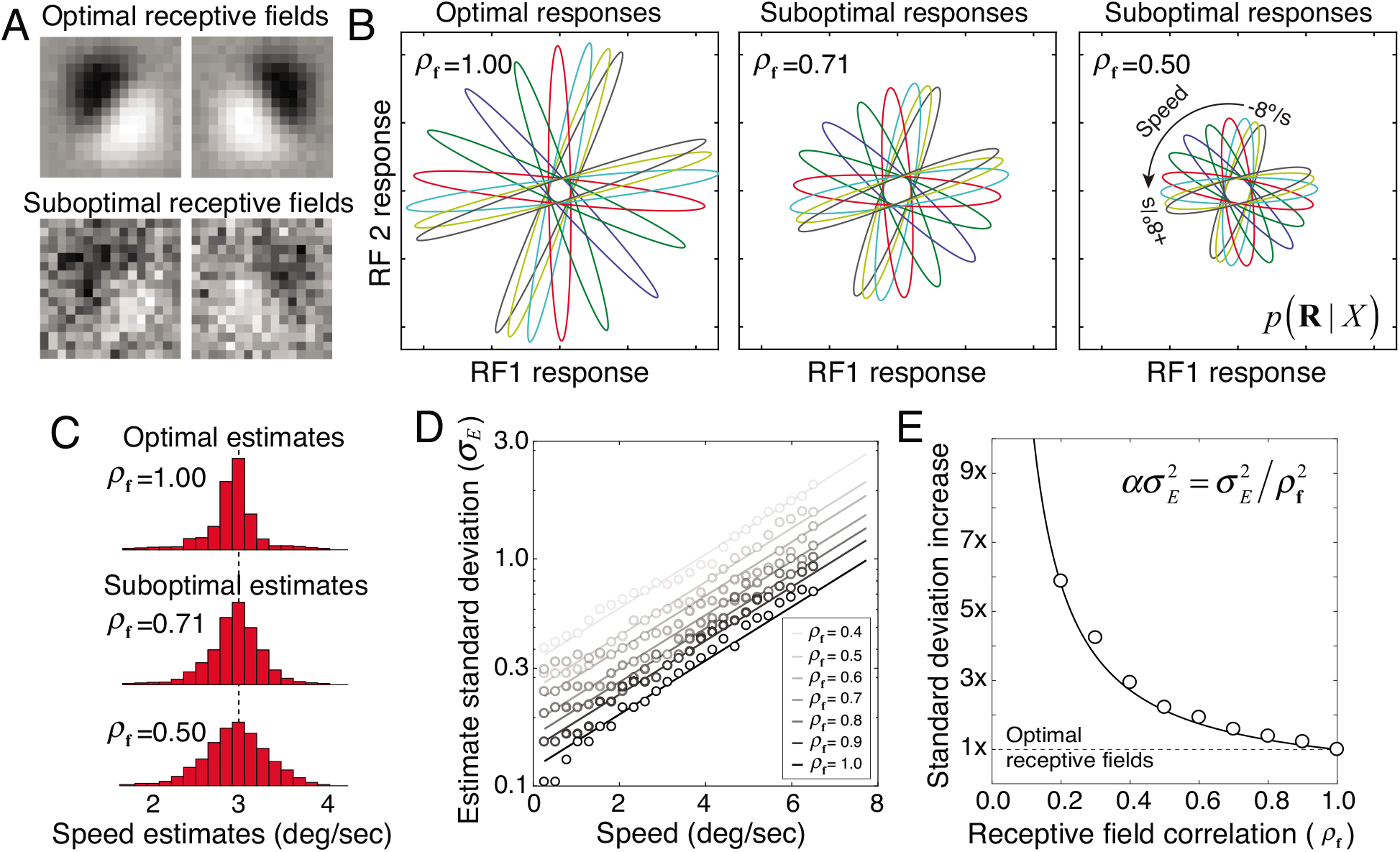
Relationship between suboptimal receptive fields and stimulus-driven variability in degraded observers. **A** Optimal receptive fields (top; also see Fig. S3) and suboptimal receptive fields from the degraded observer (bottom); only the first two receptive fields of each observer are shown. To obtain a suboptimal receptive field with a particular receptive field correlation, we added fixed samples of Gaussian white noise to the corresponding optimal receptive field. The standard deviation of the corrupting noise is given by 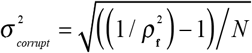 where *N* is the number of pixels defining each receptive field. **B** Impact of suboptimal receptive fields on the conditional response distributions *p*(**R**|*X*). As the receptive fields become more suboptimal, the response distributions (colored ellipses) more poorly distinguish different values of the latent variable. **C** Effect of suboptimal receptive fields on degraded observer speed estimates for movies drifting at one speed (3 deg/sec). As receptive field correlation decreases, the variance of the estimates increases, because informative stimulus features are not encoded and uninformative features are. **D** Standard deviation of degraded ideal observer speed estimates for all speeds, for a range of receptive field correlations (gray levels). Standard deviation increases exponentially with speed, which means that the standard deviations are fit by a line in log-linear space *σ_E_* = exp(*mX + b*) = exp(*b*)exp(*mX*) where *b* is the y-intercept and *m* is the slope. Note that the intercept *b*(*ρ*) depends on receptive field correlation whereas the slope does not. **E** The ratio of standard deviations for the degraded vs. the ideal observer estimates, assuming that the degraded observer has no late internal noise. The proportional increase is defined by 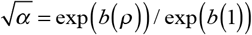 where *b*(1) is the y-intercept in D for the optimal receptive fields. Symbols plot the proportional increase from the simulations. The curve shows the expected proportional increase assuming that the scale factor is given by *α* = 1/*ρ*^2^, the relationship described in the main text. The stimulus-driven variance of the speed estimates is scaled by the squared inverse of receptive field correlation.

**Figure S10.**
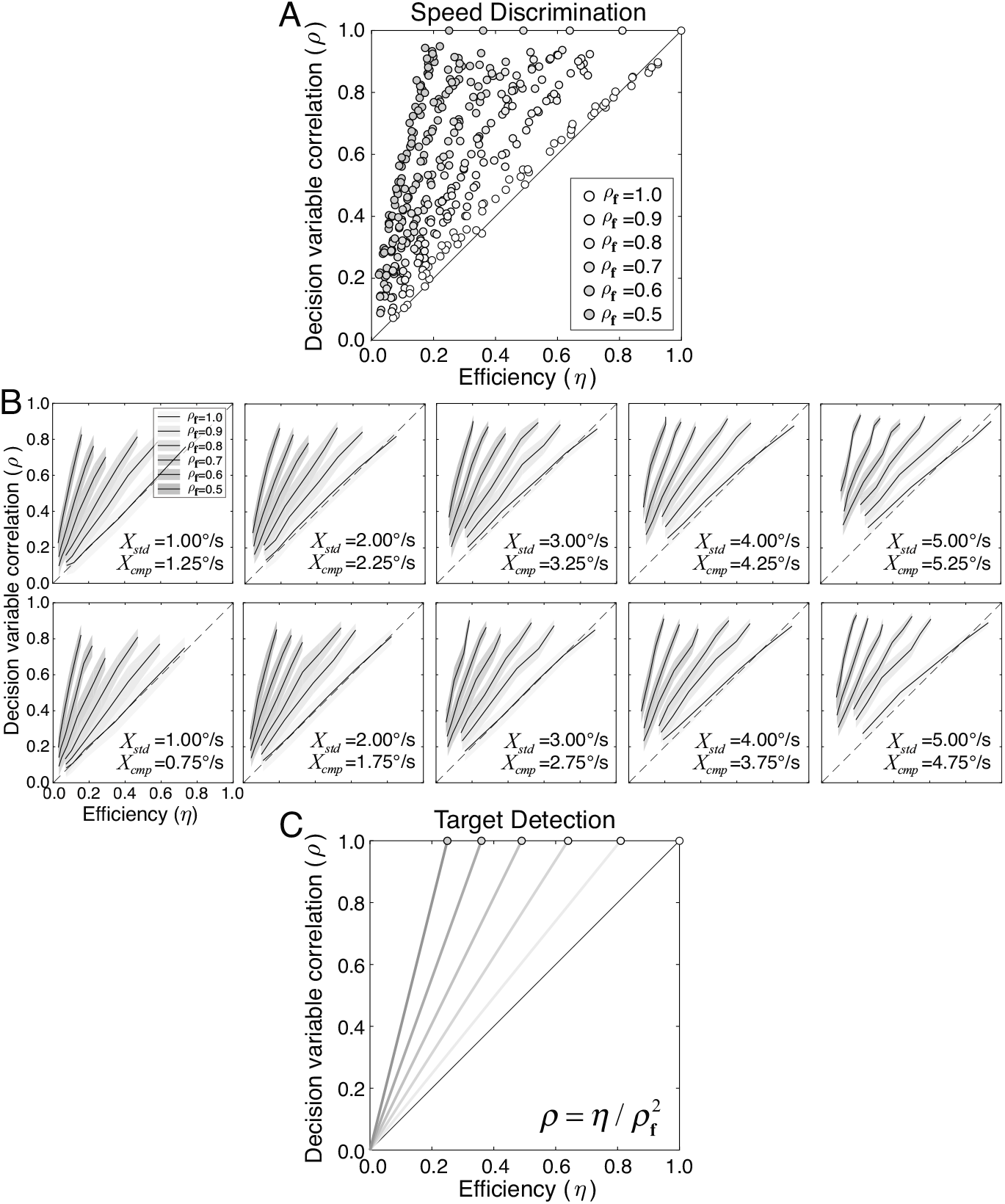
Decision variable correlation vs. efficiency for degraded observers. **A** Relationship between decision variable correlation and efficiency for degraded observers different combinations of fixed sub-optimal computations and internal noise (colors). Points represent mean decision variable correlation and mean efficiency from 100 Monte Carlo simulations of each degraded observer. White points correspond to the original ideal observer. Colored points correspond to degraded observers with receptive field correlations less than 1.0. With suboptimal computations, the decision variable correlation is proportional to efficiency with a constant of proportionality 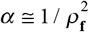 that is approximately equal to the squared inverse of receptive field correlation: for example, if receptive field correlation is 0.5, decision variable correlation is 4x higher than efficiency. **B** The relationship holds for many different combinations of standard and comparison speeds (subpanels; columns and rows represent different standard speeds and comparison speed increments, respectively; all standard and comparison speeds yield similar results.). To simulate performance for a given degraded observer we first randomly selected 1000 pairs of stimuli, each pair consisting of a standard stimulus and a comparison stimulus. We then generated speed estimates for each stimulus in the pair, subtracted the estimates to obtain a difference signal 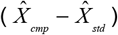, and added Gaussian noise to each difference signal to obtain the decision variable. The observer chose the comparison as faster if the decision variable exceeded zero. Efficiency and decision variable correlation were estimated from the responses of the degraded ideal observer using the same methods used for the human observers. Curves show decision variable correlation as a function of efficiency for different receptive field correlations. Shaded regions represent standard error of the decision variable correlation calculated from 100 Monte Carlo simulations. Averaging the results across conditions is therefore justified. **C** The relationship between decision variable correlation and efficiency (i.e. 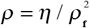) can be solved analytically for the task of target detection in Gaussian white noise (see Supplement). Interestingly, the same relationship between decision variable correlation and efficiency holds for target detection in white noise and for speed discrimination with naturalistic image movies, two tasks with very different computational requirements.

**Figure S11.**
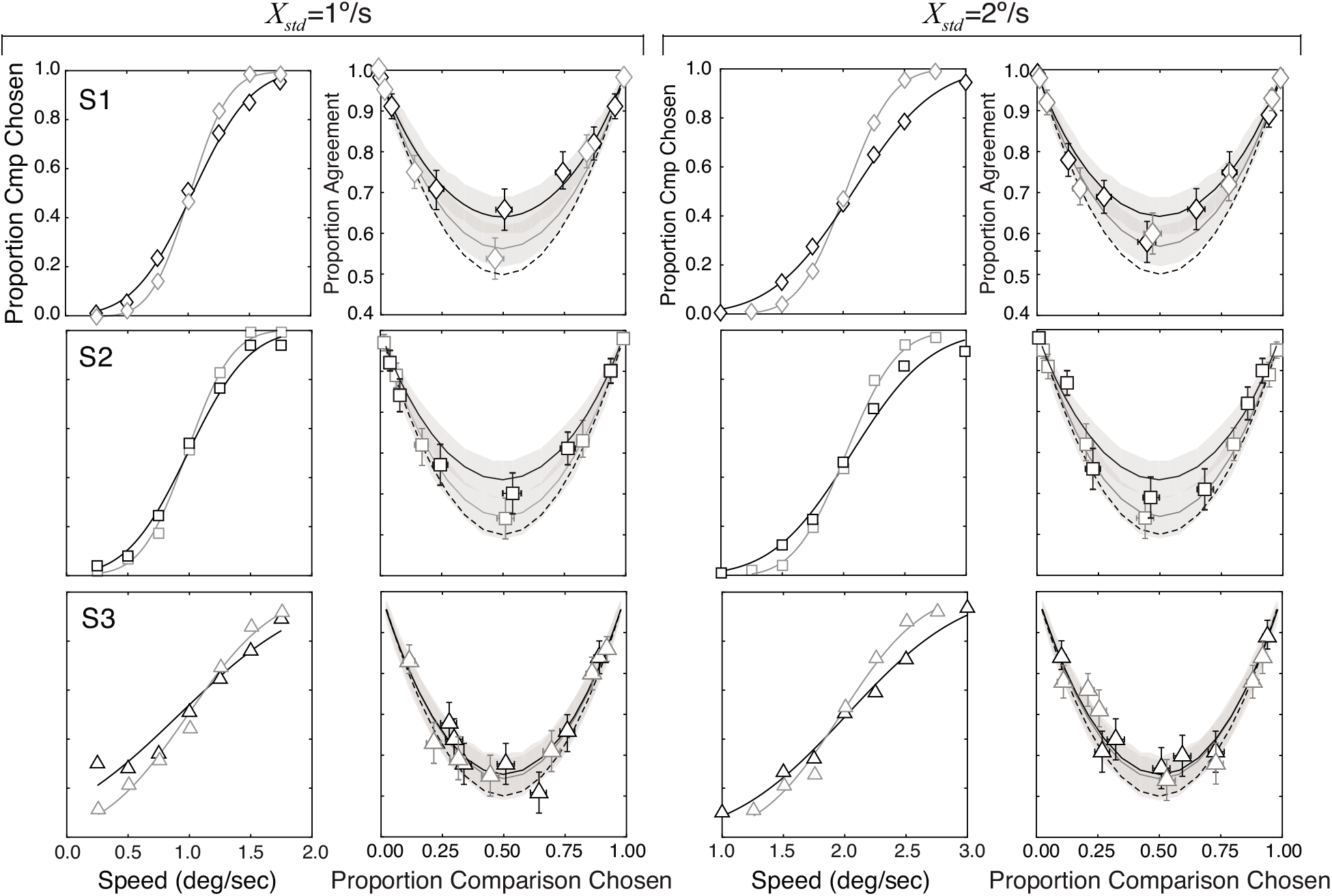
Effects of reducing stimulus variability. First and third columns show psychometric functions for speed discrimination for each human observer with drifting sinewave stimuli (gray curve) and naturalistic stimuli (black curve), with 1.0 deg/sec and 2.0 deg/sec standard speeds. Speed can be discriminated more precisely with sinewave stimuli than with naturalistic stimuli. Second and fourth columns show response agreement vs. proportion comparison chosen with sinewave stimuli (gray) and natural stimuli (black), for each human observer.

#### Efficiency vs. decision variable correlation for target detection in noise

The ideal observer for detecting a known target in Gaussian white noise requires a single receptive field that is shaped exactly like the target. If a human observer is modeled to use the exact same receptive field as the ideal observer but has a source of internal noise, the relationship between human efficiency and decision variable correlation can be determined analytically. If a human observer is modeled as using a suboptimal receptive field in addition to internal noise, decision variable correlation is proportional to efficiency scaled by the square of the receptive field correlation. Thus, even though target detection in Gaussian white noise is a significantly less complicated computational task than speed discrimination with natural images, the same equations relate efficiency and decision variable correlation.

#### Proof

Ideal observer (M) and human (H) receptive fields

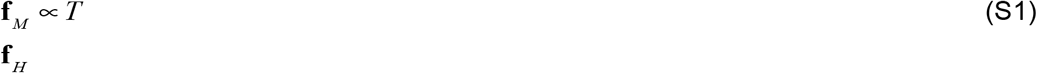

Model and human decision variables

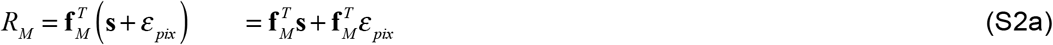

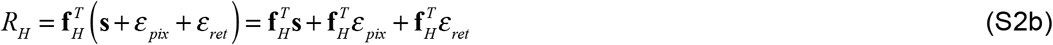

where **S** is an arbitrary stimulus, 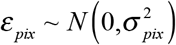 is pixel noise added to the stimulus, and 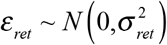 is retinal noise internal to the human observer.

Sensitivity (i.e. d-prime) is quantified by

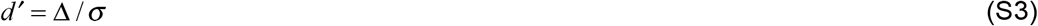

where Δ is the mean decision variable when the signal plus noise is presented and *σ* is the standard deviation of the decision variable. (The mean decision variable equals 0.0 when only noise is presented.)

The means of the model and human decision variable are given by

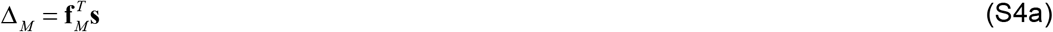

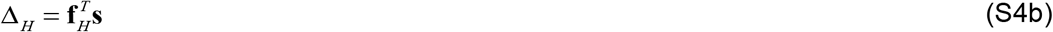

The standard deviations of the model and human decision variable are given by

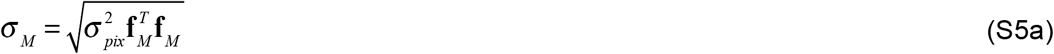

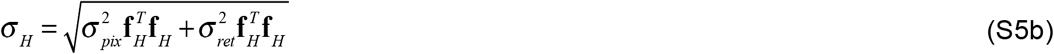

Human efficiency is the squared ratio of human and ideal sensitivity

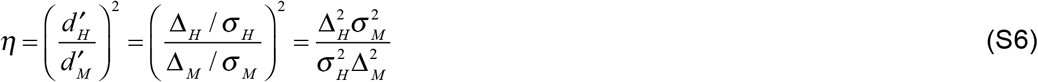

Plugging Eqs. 4 and 5 into Eq. 6 yields

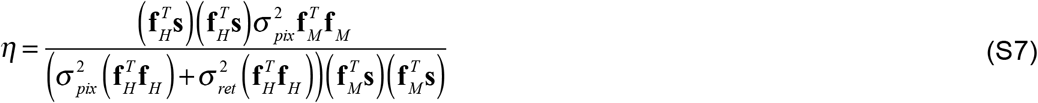

Grouping terms and simplifying

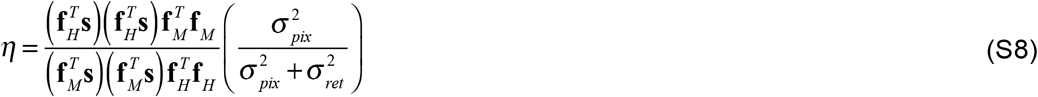

Assuming, as is typical in a target detection experiment, that the noiseless target stimulus is proportional to the target stimulus **s** = *a***f**_*M*_ yields

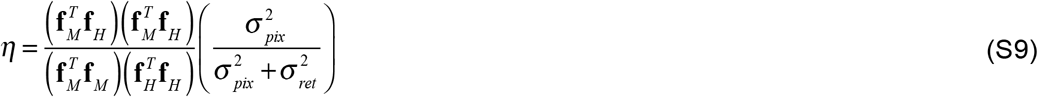

The human and ideal receptive field correlation (i.e. cosine similarity) can be used to quantify the optimality or sub-optimality of the human computations

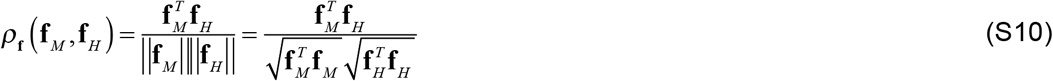

Human efficiency can be expressed in terms of receptive field correlation, internal noise, and external noise by plugging the square of Eq. S10 into Eq. S9

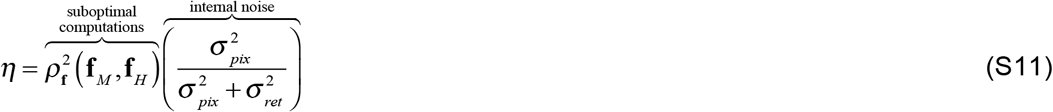

Thus, sources of inefficiency in target detection can be partitioned into fixed sub-optimal computations and internal noise. Note that if the human uses the optimal computations (i.e. uses the same receptive field **f**_*H*_ = **f**_*M*_ as the ideal observer) receptive field correlation equals 1.0 and Eq. S11 reduces to

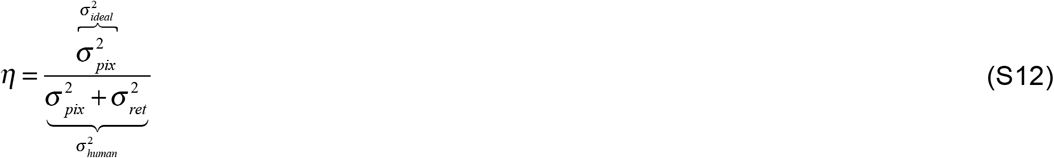

Substituting 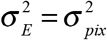 for the variance of the ideal decision variable and 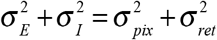 for the variance of the human decision variable into Eq. S12 yields the expression in Eq. 1 in the main text.

Next, we solve for human decision variable correlation in a double-pass target detection experiment. In a double pass experiment, the sample of external noise *ε_pix_* is identical (i.e. perfectly correlated) on repeated presentations of the same trial. From the definition of correlation, human decision variable correlation is given by

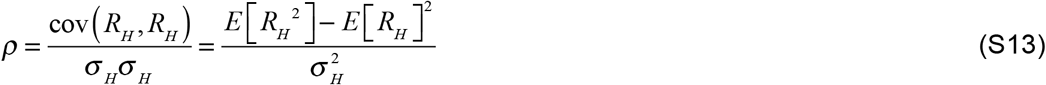

Plugging Eqs. S2b and S5b into Eq. S13, expanding, canceling terms w. independent noise across passes, and finding the variance of terms with correlated noise across passes yields

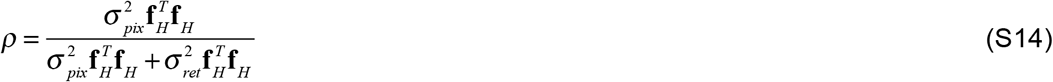

Assuming that the human uses the same receptive field across both passes of a double pass target detection experiment

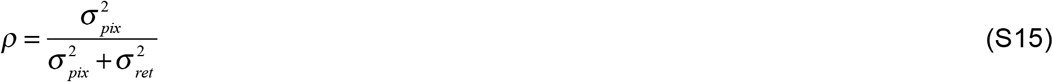

Substituting Eq. S15 into Eq. S11 and rearranging yields human decision variable correlation in terms of efficiency and receptive field correlation

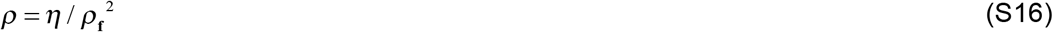

which is the same expression as Eq. 4 in the main text. Thus, decision variable correlation equals efficiency only when noise is the sole source of inefficiency, under the assumption that the human uses the same receptive field on both passes.

The expressions relating efficiency and decision variable correlation for the task of target detection in Gaussian white noise which we have just proved analytically, and in speed discrimination with naturalistic image movies which we have earlier shown via simulation (see Fig. S10), are identical.

